# *Drosophila* Fezf coordinates laminar-specific connectivity through cell-intrinsic and cell-extrinsic mechanisms

**DOI:** 10.1101/231399

**Authors:** Ivan J. Santiago, Jing Peng, Curie Ahn, Burak Gür, Katja Sporar, Chundi Xu, Aziz Karakhanyan, Marion Silies, Matthew Y. Pecot

## Abstract

Laminar arrangement of neural connections is a fundamental feature of neural circuit organization. Identifying mechanisms that coordinate neural connections within correct layers is thus vital for understanding how neural circuits are assembled. In the medulla of the *Drosophila* visual system neurons form connections within ten parallel layers. The M3 layer receives input from two neuron types that sequentially innervate M3 during development. Here we show that M3-specific innervation by both neurons is coordinated by *Drosophila* Fezf (dFezf), a conserved transcription factor that is selectively expressed by the earlier targeting input neuron. In this cell, dFezf instructs layer specificity and activates the expression of a secreted molecule (Netrin) that regulates the layer specificity of the other input neuron. We propose that employment of transcriptional modules that cell-intrinsically target neurons to specific layers, and cell-extrinsically recruit other neurons is a general mechanism for building layered networks of neural connections.

## INTRODUCTION

Precise neural connectivity underlies the structural organization and function of the nervous system. In the nervous systems of diverse organisms neural connections are arranged into layered networks, wherein each layer is defined by a unique cellular composition and the axonal arborizations of different input neurons are restricted to specific layers. This organizational strategy is thought to optimize the precision of synaptic connectivity and information processing.Layered connections define many regions of the vertebrate nervous system, including the cerebral cortex (mammals), the spinal cord and retina, and connections in the insect optic lobe are also organized in a layer-specific manner. Elucidating how neurons organize into layered networks is crucial for understanding how the precision of neural connectivity is achieved, and may provide key insights into neural function. To address the molecular and cellular bases of layer-specific connectivity we have taken a genetic approach to study layer assembly in the *Drosophila* visual system.

In the *Drosophila* visual system (Fig. 1A), photoreceptors (R1-R8) in the retina transmit signals that converge on the medulla neuropil of the optic lobe, wherein they are processed within parallel layers. The medulla receives input directly from R7 and R8 photoreceptors, which are UV and blue/green sensitive, respectively, and indirectly from broadly tuned photoreceptors (R1-R6) through second order lamina monopolar neurons L1-L5 (lamina neurons). Photoreceptors and lamina neurons project axons that innervate discrete layers and synapse with specific types of medulla neurons that process information and transmit signals to higher centers. Information from specific regions of the visual field is processed in modular columnar units, arranged perpendicular to the layers. Lamina and photoreceptor axons carrying input from the same point in space converge on targets in the same column. Input from neighboring points in space is processed in neighboring columns, establishing a retinotopic map.

**Figure 1.**
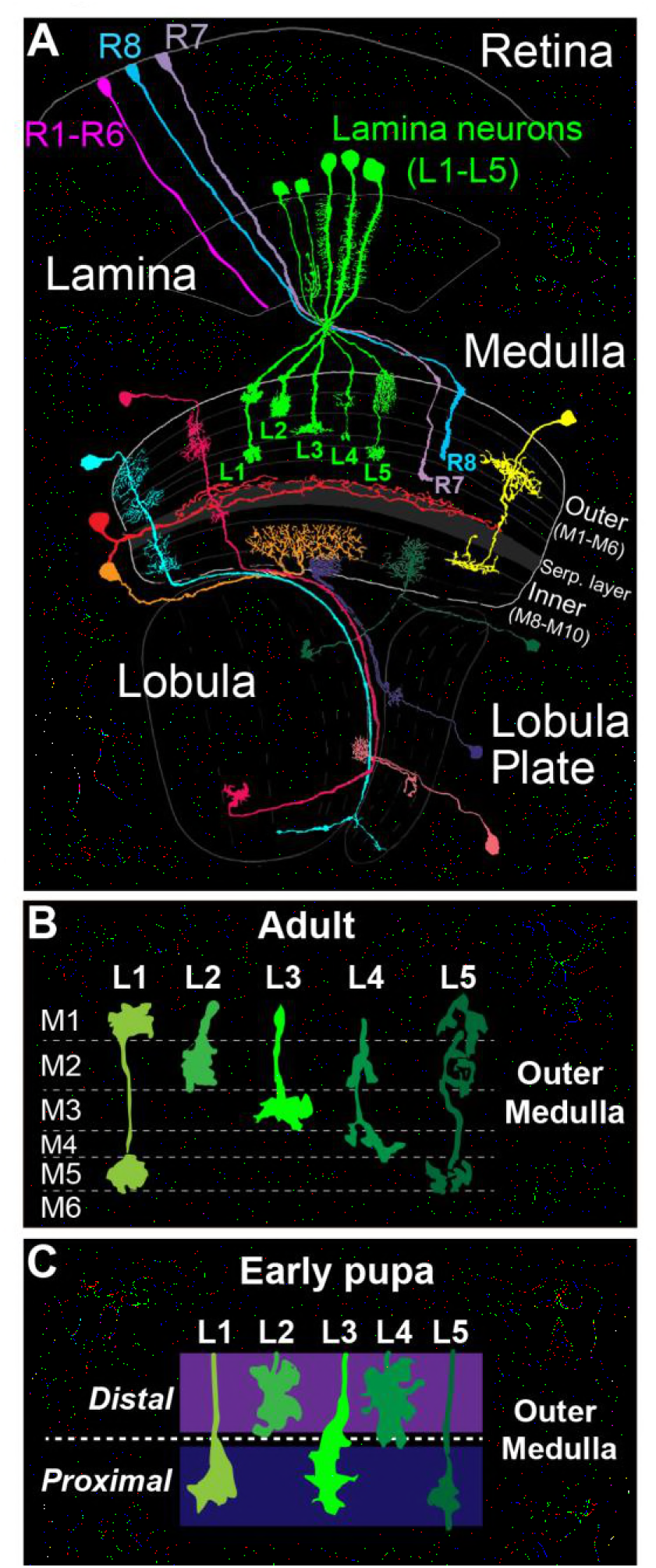
The *Drosophila* visual system and lamina monopolar neurons. (A) Anatomy of the *Drosophila* visual system (Adapted from Fischbach and Dittrich 1989). The optic lobe comprises four consecutive neuropil regions called the lamina, medulla, lobula and lobula plate. (B) Cartoon of lamina neuron axons in adult flies. The nearly mutually exclusive axonal arborizations of lamina neurons help define layers M1-M5. (C) Cartoon of lamina neuron growth cones in early pupal development. Prior to innervating discrete layers, lamina growth cones terminate in broad distal or proximal domains within the outer medulla.

The medulla comprises ten layers (M1-M10) organized into outer (M1-M6) and inner (M8-M10) regions that are divided by tangentially projecting processes that form the serpentine layer (i.e. M7) (Fischbach and Dittrich, 1989) (Fig. 1A). The cell bodies of medulla neurons are excluded from the layers, and thus layered connections develop within a dense meshwork of cellular processes. Medulla layers are defined in adult flies by the morphologies of the axon terminals and dendritic branches of particular cell types, which in general overlap completely or not at all (Fig. 1B). The positions of these processes are largely indicative of the location of synapses, although some neurites do not stratify and form synapses en passant (Takemura et al., 2013; Takemura et al., 2008; Takemura et al., 2015). In total, ~40,000 neurons that fall into more than sixty cell types form connections within one or more layers (Fischbach and Dittrich, 1989).

Studies of lamina neuron and photoreceptor axon development indicate that medulla layers emerge dynamically from broad domains. The morphologies of L1-L5 axons define layers M1- M5 in the outer medulla of adult animals (Fischbach and Dittrich, 1989) (Fig. 1B). However, in early pupal development the outer medulla is a fraction of its adult size, and lamina growth cones terminate in two broad domains (Fig. 1C) (Nern et al., 2008; Pecot et al., 2013). L2 and L4 growth cones terminate in a distal domain of the outer medulla, while L1, L3 and L5 growth cones terminate in a proximal domain. The mechanisms underlying specificity for the distal or proximal domain are not known. Nevertheless, these findings indicate that lamina neurons innervate broad domains prior to segregating into discrete layers.

Characterization of the targeting of L3 axons and the axons of R8 photoreceptors to the M3 layer has provided insight into how discrete layers emerge from broad domains. L3 and R8 axons innervate M3 sequentially during development. R8 neurons are born first and project axons that terminate near the distal surface of the medulla (known as M0), where they remain for up to two days (Ting et al., 2005). L3 axons within the same column project past R8 axons and terminate within a broad domain in the proximal outer medulla (discussed above) (Nern et al., 2008; Pecot et al., 2013). At this stage L3 growth cones are broad, nearly spanning the outer medulla. Subsequently, L3 growth cones segregate into the developing M3 layer through a stereotyped structural rearrangement that is regulated by mechanisms that are not well understood (Pecot et al., 2013). Within M3, L3 growth cones secrete Netrin, which acts locally to regulate the attachment of R8 growth cones within the layer after they extend from M0 (Akin and Zipursky, 2016; Pecot et al., 2014; Timofeev et al., 2012). Thus, the M3 layer develops in part through the stepwise innervation of L3 and R8 axons, and R8 layer specificity depends on a signal from L3 growth cones.

Taken together, these developmental studies point to a model, wherein outer medulla layers emerge in a stepwise manner from broad domains through a precise sequence of interactions between specific cell types. However, while molecules that are necessary for layer specificity in isolated contexts have been identified, whether overarching mechanisms coordinate the assembly of specific layers is unclear. To address this, we concentrate on assembly of the M3 layer, and in this study focus on L3 neurons, for which we have developed an assortment of genetic tools. Previously, we found that *Drosophila* Fezf (dFezf)/Earmuff, the *Drosophila* ortholog of the Fezf zinc finger transcription factors in vertebrates (Fezf1/2) (Hashimoto et al., 2000a; Hashimoto et al., 2000b; Matsuo-Takasaki et al., 2000; Pfeiffer et al., 2008; Weng et al., 2010), is selectively expressed in L3 neurons in the lamina (Tan et al., 2015). Thus, we reasoned that dFezf might regulate the expression of genes that control cell-specific aspects of L3 development, including broad domain specificity, layer specificity, and synaptic specificity within the target layer.

Interestingly, our genetic analyses of dFezf function in L3 neurons revealed a role for dFezf in regulating the targeting of L3 *and* R8 axons within the medulla. DFezf functions cell autonomously to promote the targeting of L3 growth cones to the proximal domain of the outer medulla, which is essential for L3 layer specificity. We also find that dFezf non-autonomously regulates R8 layer specificity through the activation of *Netrin* expression in L3 neurons. When dFezf function is lost in L3 neurons, L3 and R8 axons innervate inappropriate layers while the dendrites of a common synaptic target (Tm9) innervate the M3 layer normally. As a result, synaptic connectivity with the target cell is disrupted. We conclude that dFezf represents a transcriptional module that constructs M3 layer circuitry by controlling the layer innervation of specific cell types through cell-intrinsic and cell-extrinsic mechanisms. We propose that the use of such modules to coordinate neural connections within correct layers is a widespread strategy for building laminar-specific circuits.

## RESULTS

### DFezf is selectively expressed in L3 neurons during pupal development and in adult flies

As a first step towards characterizing the function of dFezf in L3 neurons we examined its expression during development. Previously, we showed that dFezf is expressed in L3 neurons at 40 hours after puparium formation (h APF) (Tan et al., 2015), when L3 growth cones are in the process of segregating into the developing M3 layer (Pecot et al., 2013). To assess if dFezf is also expressed in L3 neurons at other stages of development and determine whether expression remains restricted to L3 neurons, we assessed dFezf immunostaining in the lamina using confocal microscopy. We discovered that dFezf is expressed in L3 neurons from early pupal stages (at least as early as 12h APF) through to adulthood, but is not expressed in late third instar larvae when lamina neurons begin to differentiate (Fig. 2A-D). DFezf expression was not detected in lamina neuron precursor cells or any other cells besides L3 at any stage of development in the lamina. The timing and selectivity of dFezf expression is consistent with dFezf playing an important role in regulating L3 development.

**Figure 2.**
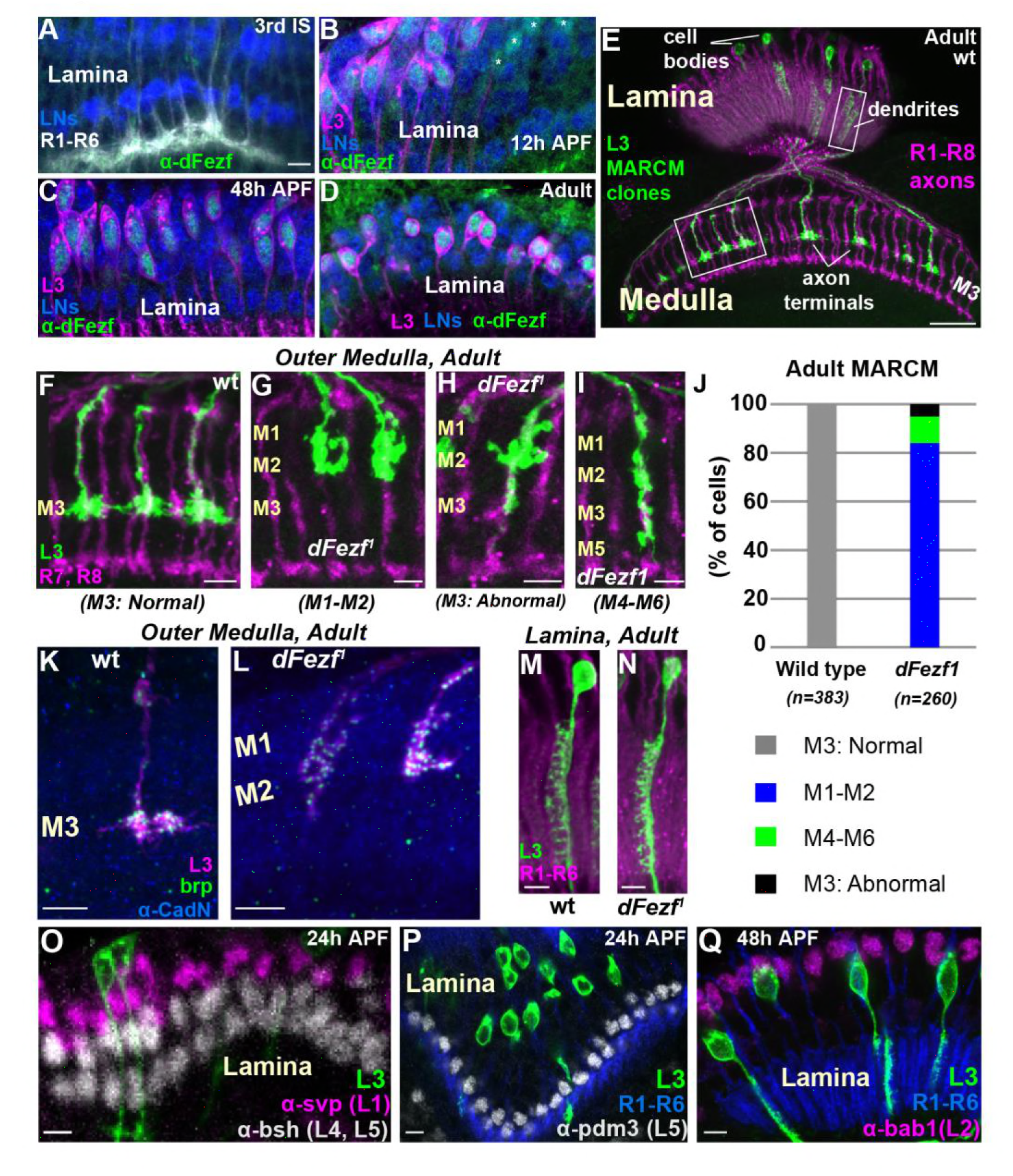
*DFezf* is cell autonomously required for L3 layer specificity and axon terminal morphology. (A-D) Confocal images showing dFezf protein expression in the lamina from the late 3^rd^ Instar larval stage through to adulthood (newly eclosed flies). DFezf expression (green) was observed through immunohistochemistry using a specific antibody. Elav immunostaining (blue) labels all lamina neurons (LNs). At least five brains were examined per time point. Scale bar = 5 microns and applies to all images. (A) DFezf (green) is not expressed in the lamina during the 3^rd^ Instar larval stage. Lamina neurons (blue) differentiate in close proximity to R1-R6 axons (white, mAB24B10) forming columnar-like cartridges oriented orthogonally to the lamina plexus (thick white band) comprising R1-R6 terminals. (B-D) DFezf (green) is expressed exclusively in L3 neurons in the lamina during pupal development and in newly eclosed adults. L3 neurons (magenta) express myr-GFP (anti-GFP) driven by 9-9-GAL4 (Nern et al., 2008). Asterisks in (B) indicate dFezf-expressing lamina neurons (green + blue) that are most likely L3 neurons that have yet to turn on GAL4 expression (youngest L3 neurons). (E-N) MARCM + STaR experiments in adult flies. See methods section for a detailed description. Scale bars when shown are five microns with the exception of the scale bar shown in (E), which is twenty microns. L3 clones (green) expressed myr-tandem Tomato (tdTOM) and were visualized using a DsRed antibody. (E) Broad view (confocal image) of the morphologies of wild type L3 clones (green) in the lamina and medulla, which are indicated by R1-R8 axons (magenta, mAB24B10). Boxed regions indicate examples of the regions shown in confocal images in F-I, M and N. (F-I) Confocal images of terminals from wild type or *dFezf^1^* L3 neurons (green), within the outer medulla (M1-M6), defined by R7 and R8 axons (magenta, mAB24B10). (J) Adult MARCM quantification. N is the total number of neurons counted per genotype. 10 brains were analyzed for each genotype. (K, L) Confocal images show L3 clones (magenta) within the outer medulla and, within L3 terminals, Brp (green) expression and localization. Brp expressed from its native promoter within a bacterial artificial chromosome (BAC) was selectively tagged with smFPV5 in L3 MARCM clones, and visualized using an anti-V5 antibody. N-Cadherin (CadN) immunostaining (blue) serves as a neuropil marker. All *dFezf^1^* L3 neurons displayed Brp puncta in their terminals. At least five brains were analyzed for each genotype. (M and N) Morphologies of dendrites from wild type or *dFezf^1^* L3 neurons in the lamina. The lamina is indicated by R1-R6 axons (magenta, mAB24B10) (O-Q) MARCM experiments analyzed at 24 or 48h APF. *DFezf^1^* L3 clones (green) express myr-GFP (9-9-GAL4) and are visualized using a GFP antibody. Co-labeling of L3 somas and transcription factors specific for other lamina neuron subtypes (using specific antibodies) show that the mutant neurons do not express these proteins. (P and Q) R1-R6 axons (blue, mAB24B10) demarcate lamina cartridges and the lamina plexus.

### DFezf is cell autonomously required for L3 layer specificity

To investigate the function of *dFezf* in L3 neurons we used mosaic analysis with a repressible cell marker (MARCM) (Lee and Luo, 1999) in conjunction with synaptic tagging with recombination (STaR) (Chen et al., 2014). This allowed us to generate single, fluorescently labeled wild type or *dFezf* mutant L3 neurons while simultaneously visualizing the active zone protein Bruchpilot (Brp), expressed from its native promoter (see methods for detailed description). The *dFezf* null allele used in these experiments (*dFezf^1^*) contains a single A to T nucleotide change leading to the substitution of a leucine for a conserved histidine in the third zinc-finger domain (Weng et al., 2010). We found that L3 neurons homozygous for *dFezf^1^* displayed defects in axon terminal morphology and layer specificity in the medulla. Wild type L3 neurons elaborate stratified axon terminals within the M3 layer (100%) (Fig. 2E and F, J). The large majority (84%) of the axons of *dFezf^1^* L3 neurons terminated inappropriately within layers distal to M3 (i.e. M1-M2) (Fig. 2G, J) and had abnormally shaped terminals (compare Fig. 2F and G). The remaining 16% of the axons either terminated in the M3 layer (Fig. 2H, J) or in more proximal layers (e.g. M4-M6) (Fig. 2I, J), and these also had terminal-like swellings in M1- M2 (Fig. 2H and I). Presynaptic Brp puncta were present in mis-targeting L3 terminals (Fig. 2K and L), suggesting that mutant neurons formed synaptic connections within incorrect layers. In the lamina, L3 neurons display a unique dendritic projection pattern in which all dendrites project to one side of the neurite (Fig. 2M). Dendritic processes were largely normal in *dFezf* null neurons (compare Fig. 2M and N). In addition, the mutant L3 neurons did not express transcription factors specific to other lamina neurons (Fig. 2O-Q) and still expressed L3-specific markers early in development (see Fig. 3), showing that disrupting *dFezf* did not cause an obvious change in cell fate. However, at least some L3-specific markers turn down in *dFezf^1^* L3 neurons during late pupal development (see below), suggesting dFezf function is required to maintain their expression. Taken together, these findings demonstrate that *dFezf* is cell autonomously required for proper axon terminal morphology and layer specificity, yet dispensable for dendrite formation.

**Figure 3.**
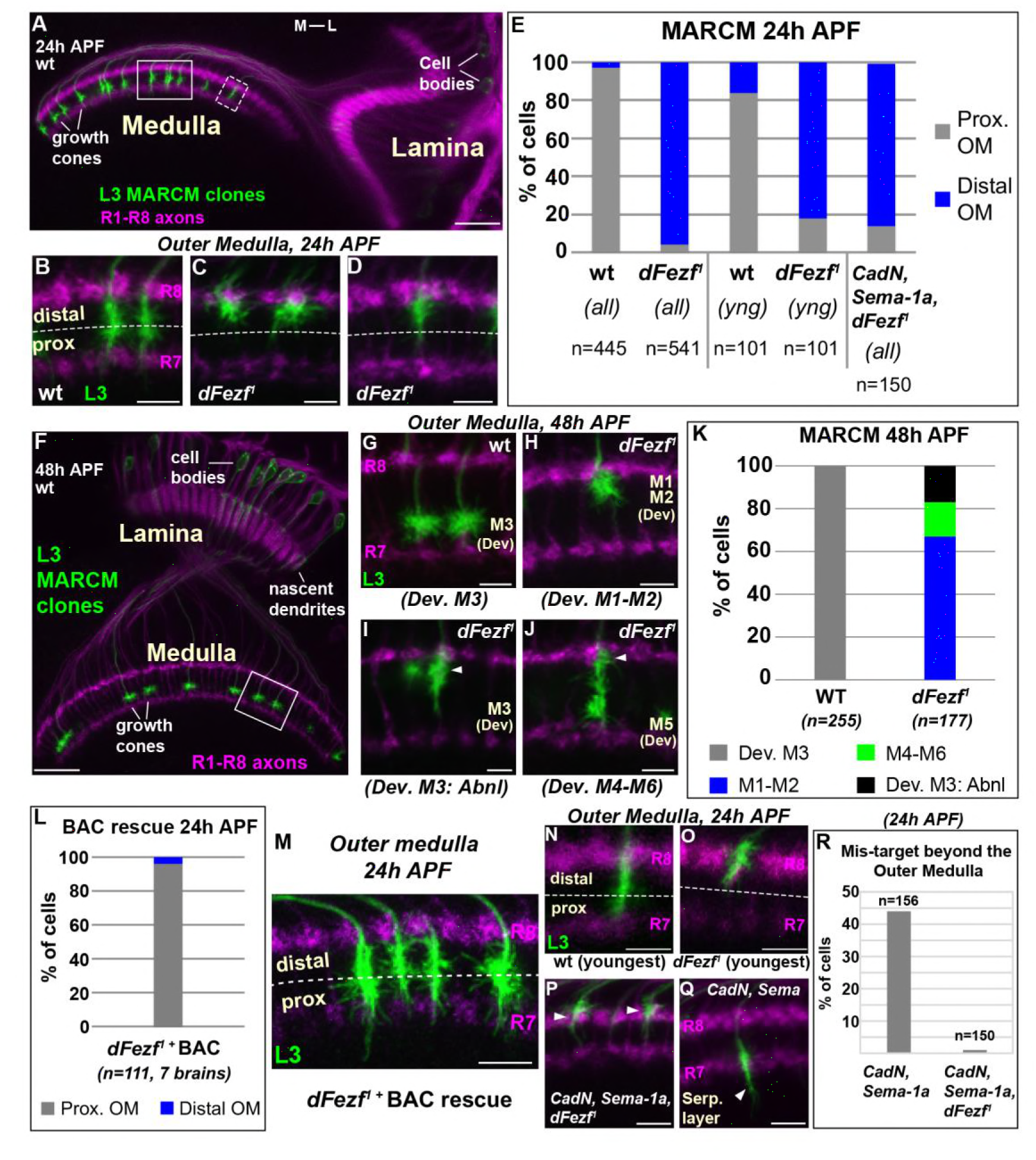
*DFezf* is required for the targeting of L3 growth cones to the proximal domain of the outer medulla. (A-R) MARCM experiments analyzed at 24 or 48h APF. L3 clones expressed myr-GFP driven by 9-9-GAL4 and were visualized in confocal images using a GFP antibody. (A) Broad view of the morphologies of wild type L3 clones (green) within the lamina and medulla (indicated by R1-R8 axons, magenta [mAB24B10]) at 24h APF. The solid boxed region indicates an example of confocal images shown in B-D. The dashed boxed region shows an example of a newly born L3 clone in the most lateral part of the medulla array similar to N and O. M-L indicates medial and lateral. Scale bar = 20 microns. (B-D, G-J, M-Q) Confocal images show a zoomed in view of L3 growth cones within the outer medulla. At these stages of development the boundaries of the outer medulla are defined by R7 and R8 growth cones (magenta, mAB24B10). Scale bars = 5 microns. (E, K, L, R) Quantification of MARCM experiments. N is the total number of neurons per genotype that were counted. (E) WT (7 brains), *dFezf*^1^ (12 brains), *CadN*, *Sema-1a*, *dFezf*^1^ (2 brains). (*all*) means that all L3 clones were counted while (*yng*) indicates that the L3 clones present within the most lateral five medulla columns containing R7 growth cones were counted (i.e. youngest L3 clones). Prox. OM= proximal domain of the outer medulla, Distal OM = distal domain of the outer medulla. (F) Broad view of the morphologies of wild type L3 clones in the lamina and medulla (indicated by R1-R8 axons, magenta [mAB24B10]) at 48h APF. The solid boxed region is an example of regions shown in confocal images in G-J. Scale bar = 20 microns. (I and J) Arrowheads indicate abnormal swellings within the developing M1-M2 layers. (K) Wild type (6 brains), *dFezf^1^* (6 brains). (L) Prox. OM = proximal domain of the outer medulla, Distal OM = distal domain of the outer medulla. (P and Q) Arrowheads indicate the depths within the medulla at which L3 growth cones terminate. The serpentine layer (indicated in Q) lies directly beneath the outer medulla (R) *CadN, Sema-1a* (2 brains), *CadN*, *Sema-1a*, *dFezf^1^* (2 brains).

### DFezf is necessary for targeting to the proximal domain of the outer medulla

To determine how disrupting *dFezf* affects L3 axon targeting during development, we compared the axonal morphologies of wild type and *dFezf^1^* neurons during pupal development using MARCM. In early pupal development (~24h APF), the axons of wild type L3 neurons terminate in the proximal domain of the outer medulla directly above R7 growth cones (Fig. 3A and B, E). The axons of *dFezf^1^* neurons incorrectly terminated within the distal domain of the outer medulla (Fig. 3 C-E). Under these conditions L3 growth cones overlapped with or terminated just underneath the growth cones of R8 photoreceptors near the distal medulla surface. By mid-pupal development (~48h APF), the growth cones of wild type L3 neurons have segregated into the developing M3 layer (Fig. 3F and G, K). However, we found that all of the growth cones of *dFezf^1^* L3 neurons displayed abnormal morphologies, defects in layer specificity, or both. Most of the abnormal growth cones (67%) exclusively innervated the developing M1 and M2 layers (Fig. 3H, K). The remaining 33% of the growth cones terminated in the developing M3 layer (Fig. 3I, K) or in more proximal layers (e.g. M4-M6) (Fig. 3J, K). However, each of these growth cones also displayed abnormal swellings in distal layers (i.e. the developing M1 and M2 layers) (Fig. 3I and J, arrowheads). These defects in morphology and layer innervation are consistent with the abnormalities observed in adult flies under these conditions. We observed similar deficits in L3 growth cone targeting in MARCM experiments when we used a different *dFezf* mutant allele (*dFezf^2^*) that eliminates the entire *dFezf* open reading frame (Weng et al., 2010) (Fig. S1). Moreover, defects in growth cone targeting during pupal development, as well as abnormal layer innervation in adult flies caused by *dFezf^1^* or *dFezf^2^* were completely rescued by a single copy of a bacterial artificial chromosome (BAC) containing the *dFezf* locus (Fig. 3L and M, Fig. S2). Together, these findings demonstrate that *dFezf* is cell-autonomously required for the targeting of L3 growth cones to the proximal domain of the outer medulla, which may be necessary for the subsequent segregation of the growth cones into the developing M3 layer.

We reasoned that the growth cones of *dFezf* null L3 neurons could directly terminate within the distal domain of the outer medulla, or could initially target correctly to the proximal domain and then retract. Direct mis-targeting would be consistent with a change in axon target specificity, while retraction would suggest a defect in stabilizing growth cone position. To distinguish between these possibilities we took two approaches. First, in MARCM experiments we focused on the L3 clones that had most recently innervated the medulla (i.e. the youngest clones, see dashed box in Fig. 3A). Unlike the youngest wild type clones, the large majority of the nascent *dFezf^1^* clones (82%) terminated within the distal domain of the outer medulla similar to their older counter parts (Fig. 3E, N and O), consistent with most of these neurons directly innervating this region. Second, we investigated growth cone targeting when *dFezf* was disrupted in combination with *CadN* and *Sema-1a*. CadN and Sema-1a function synergistically to restrict L3 growth cones to the outer medulla during pupal development, preventing innervation of the serpentine layer and inner medulla (Pecot et al., 2013). We hypothesized that if the growth cones of *dFezf^1^* L3 neurons initially target correctly and then retract, then in the absence of CadN and Sema-1a function the growth cones would inappropriately innervate the serpentine layer and inner medulla, as in *CadN* and *Sema-1a* double mutant neurons. Conversely, if the growth cones of *dFezf^1^* L3 neurons directly mis-target to the distal outer medulla, this should prevent mis-targeting beyond the outer medulla caused by the loss of CadN and Sema-1a function, and the triple KO phenotype should resemble the loss of dFezf function only. MARCM experiments revealed the latter to be correct. L3 neurons lacking the function of dFezf, CadN and Sema-1a predominantly innervated the distal domain of the outer medulla in early pupal development (Fig. 3E, P), similar to L3 neurons lacking dFezf function only (Fig. 3C-E), and did not mis-target beyond the outer medulla like *CadN/Sema-1a* double mutant L3 neurons (Fig. 3Q and R). Combined with our analyses of newly born *dFezf^1^* L3 neurons, this suggests that L3 neurons lacking dFezf function directly innervate the distal domain of the outer medulla, which is consistent with a switch in broad domain specificity.

### DFezf acts instructively to regulate growth cone targeting to the proximal domain of the outer medulla

DFezf could function permissively or in an instructive manner to regulate the targeting of L3 growth cones to the proximal domain of the outer medulla. To discriminate between these possibilities we performed mis-expression experiments in other lamina neurons. If dFezf function is instructive, then mis-expressing it in L2 or L4 neurons, which normally target to the distal domain of the outer medulla, should cause their growth cones to terminate in the proximal domain. By contrast, as L1 and L5 growth cones normally terminate in the proximal domain, dFezf-mis-expression in these neurons should not affect growth cone targeting. For technical reasons we decided to focus on L2 and L5 neurons. In L5 neurons, we mis-expressed cDNA encoding dFezf-HA using 6-60-GAL4, which has been shown to selectively drive expression in L5 neurons in the lamina (Nern et al., 2008). In these experiments all L5 neurons (and a small number of medulla neurons) expressed dFezf-HA, as determined through HA immunostaining, and a subset of these were specifically labeled by myr-GFP (Fig. 4A). We found that the expression of dFezf-HA had no effect on L5 growth cone position in early pupal development. The growth cones of dFezf-expressing L5 neurons innervated the proximal domain of the outer medulla in a manner indistinguishable from wild type neurons (Fig. 4B-D).

**Figure 4.**
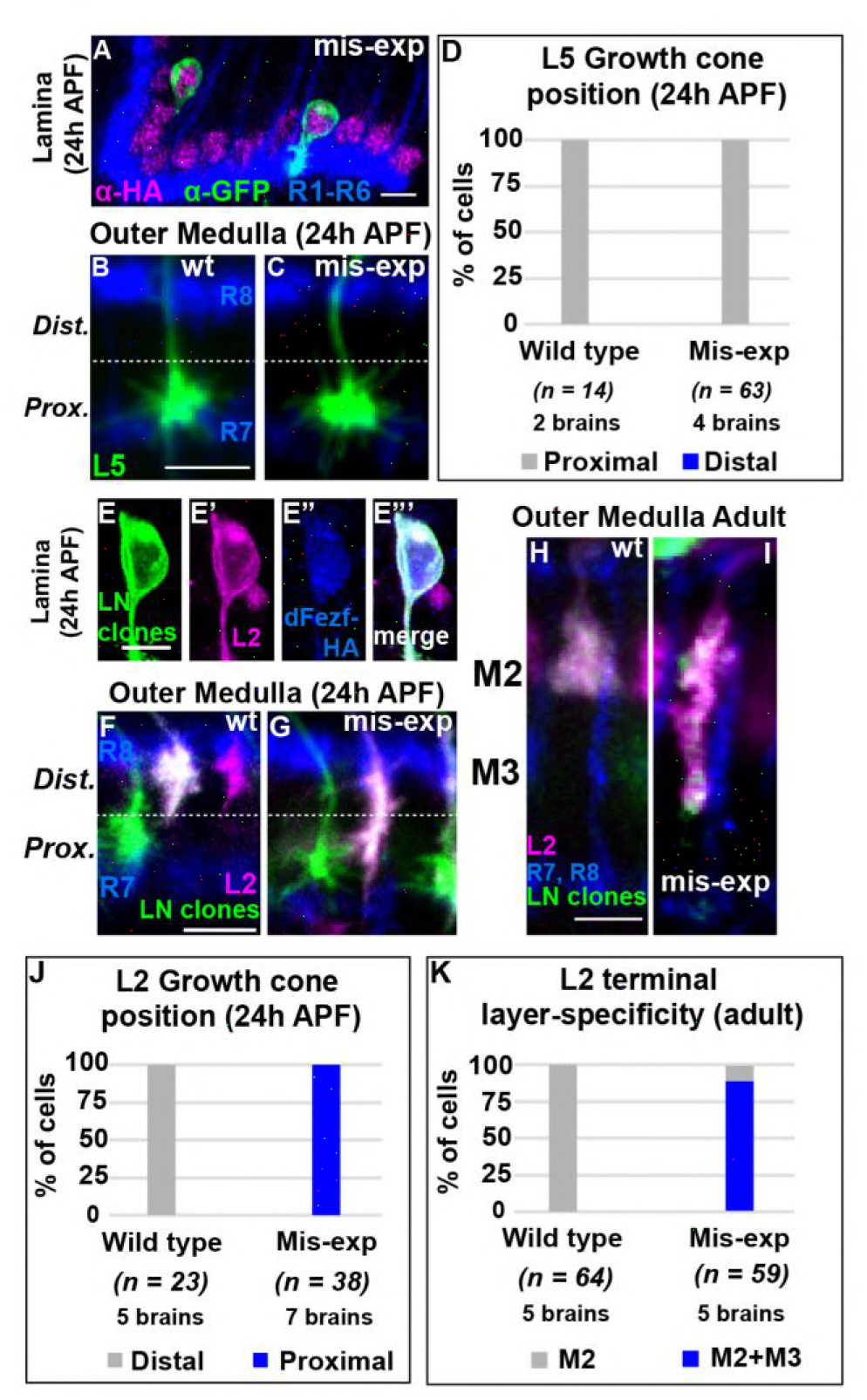
DFezf plays an instructive role in regulating growth cone targeting. (A-D) L5 mis-expression experiments. DFezf-HA was expressed in L5 neurons using 6-60- GAL4, and sparse labeling of L5 neurons (myr-GFP, α-GFP) was achieved using the FlpOut method (see methods for detailed description). (A) DFezf-HA (magenta) is expressed in all L5 neurons, a subset of which are labeled with myr-GFP (green) using FlpOut. L5 neurons are easily identified based on position, as their somas are located directly distal to the lamina plexus defined by R1-R6 axons (blue, mAb24B10), which appears as a thick band. (B and C) Confocal images of L5 growth cones (green) within the outer medulla at 24h APF. R7 and R8 growth cones (blue, mAB24B10) define the boundaries of the outer medulla. (D) Quantification of L5 mis-expression experiments. N = the number of cells counted. (E-K) Mis-expression of dFezf-HA in L2 neurons in MARCM experiments. (E-E’’’) DFezf-HA (blue, α-HA) is expressed in L2 MARCM clones, which are double positive for myr-GFP (green, 11-164-GAL4) and myr-tdTOM (magenta, 39D12-LexA). (F-I) Confocal images showing L2 targeting. Lamina neuron clones (green) were labeled with myr-GFP (α-GFP) driven by 11-164-GAL4 (Nern et al., 2008). In G and I dFezf-HA was also expressed in lamina neuron clones by 11-164-GAL4. L2 neurons (magenta) were labeled with myr-tdTOM (α-DsRed) driven by 39D12-LexA. L2 MARCM clones are double positive for myr-GFP and myr-tdTOM and appear white. Scale bars = 5 microns. (F and G) R7 and R8 growth cones (blue, mAB24B10) define the boundaries of the outer medulla at 24h APF. (H and I) R7 and R8 axons (blue, mAB24B10) define outer medulla layers in adult flies. (J and K) Quantification of L2 mis-expression experiments. N = the number of cells counted.

To assess the effects of mis-expressing *dFezf* in L2 neurons, we expressed cDNA encoding dFezf-HA in a subset of lamina neurons using a pan-lamina neuron GAL4 driver (11-164-GAL4) in MARCM experiments, and labeled L2 neurons using an L2-specific LexA driver (39D12-LexA). The expression of dFezf-HA was assessed through HA immunostaining (Fig. 4E). We found that, in early pupal development, the growth cones of all L2 neurons mis-expressing dFezf-HA inappropriately terminated within the proximal domain of the outer medulla (Fig.4F and G, J), and in adult flies the large majority of these neurons (88%) incorrectly innervated the M3 layer (Fig. 4H and I, K). In all cases dFezf-HA expressing L2 neurons still displayed terminal-like swellings within the M2 layer, indicating a partial rather than complete change in layer specificity. These experiments demonstrate that mis-expressing *dFezf* in L2 neurons is sufficient to induce innervation of the proximal domain of the outer medulla and cause ectopic innervation of the L3 target layer M3. Collectively, our mis-expression studies in L2 and L5 neurons support that dFezf acts instructively to promote growth cone targeting to the proximal domain of the outer medulla.

### DFezf functions in parallel to CadN and Sema-1a to regulate L3 growth cone targeting

CadN and Sema-1a play a crucial role in L3 growth cone targeting (Nern et al., 2008; Pecot et al., 2013), and we next sought to characterize the relationship between *dFezf, CadN* and *Sema-1a* in L3 neurons. We hypothesized that dFezf could regulate the expression of *CadN* and *Sema-1a* in L3 neurons, potentially as part of a gene program controlling growth cone targeting, or dFezf could function in parallel to CadN and Sema-1a. If the former is true, then disrupting *dFezf* in L3 neurons should alter *CadN* and *Sema-1a* expression. Transcriptional profiling of L3 MARCM clones at 40h APF using RNA-seq showed that the levels of *CadN* and *Sema-1a* mRNA were not significantly changed when *dFezf* was disrupted (Fig. S3A and B) (the remaining RNA-seq data will be published elsewhere). These findings show that *CadN* and *Sema-1a* expression in L3 neurons does not require *dFezf*. In further support of this, we found that while *CadN* and *Sema-1a* double mutant L3 neurons display defects in dendrite morphology (Fig. S3C), dendrites are qualitatively normal in the absence of dFezf function (Fig. 2M and N), indicating that disrupting *dFezf* does not affect CadN and Sema-1a function. Together, these data indicate that dFezf acts in parallel to CadN and Sema-1a to regulate L3 growth cone targeting within the outer medulla (see discussion).

### *DFezf is necessary and sufficient for* Netrin *expression*

Our RNA-seq experiments revealed that the mRNA levels of *Netrin-A* and *B* (*Net-A/B*) were drastically reduced in *dFezf* null L3 neurons compared to wild type neurons at 40h APF (Fig. S3A and B), indicating these genes are direct or indirect dFezf targets. This suggested that, in addition to cell autonomously regulating L3 targeting, dFezf non-autonomously regulates R8 layer specificity through the activation of *Netrin* in L3 neurons. To test this we first assessed Netrin protein expression in L3 neurons during development. Previous gene expression and genetic studies implied that L3 neurons express Netrin protein (Pecot et al., 2014; Tan et al., 2015; Timofeev et al., 2012), but this had not been shown directly. To address this we used a Net-B-GFP protein trap, wherein endogenous Net-B is tagged with GFP. At 24h APF we observed Net-B expression in the perinuclear region of the oldest lamina neurons (most anterior) (Fig. 5A-D’). Using markers specific for different lamina neuron subclasses we discovered that Net-B is expressed by L3, L4 and L5 neurons, but not L1 or L2 neurons. We assessed Net-B expression in L3 by labeling L3 somata with FLAG-tagged dFezf (dFezf-FLAG) expressed from its native promoter within a BAC (Fig. S4). At 48h APF Net-B expression remained specific to L3, L4 and L5 neurons in the lamina, although it was most strongly expressed in L3 neurons at this stage (Fig. 5E-H). Thus, while Netrin immunoreactivity is first detected within the M3 layer at ~40h APF (Timofeev et al., 2012), which coincides with the segregation of L3 growth cones into the developing M3 layer, L3 neurons express Netrin much earlier in development. In addition, we found that Netrin is also expressed by L4 and L5 neurons during pupal development, and thus may regulate circuit assembly within multiple developing layers.

**Figure 5.**
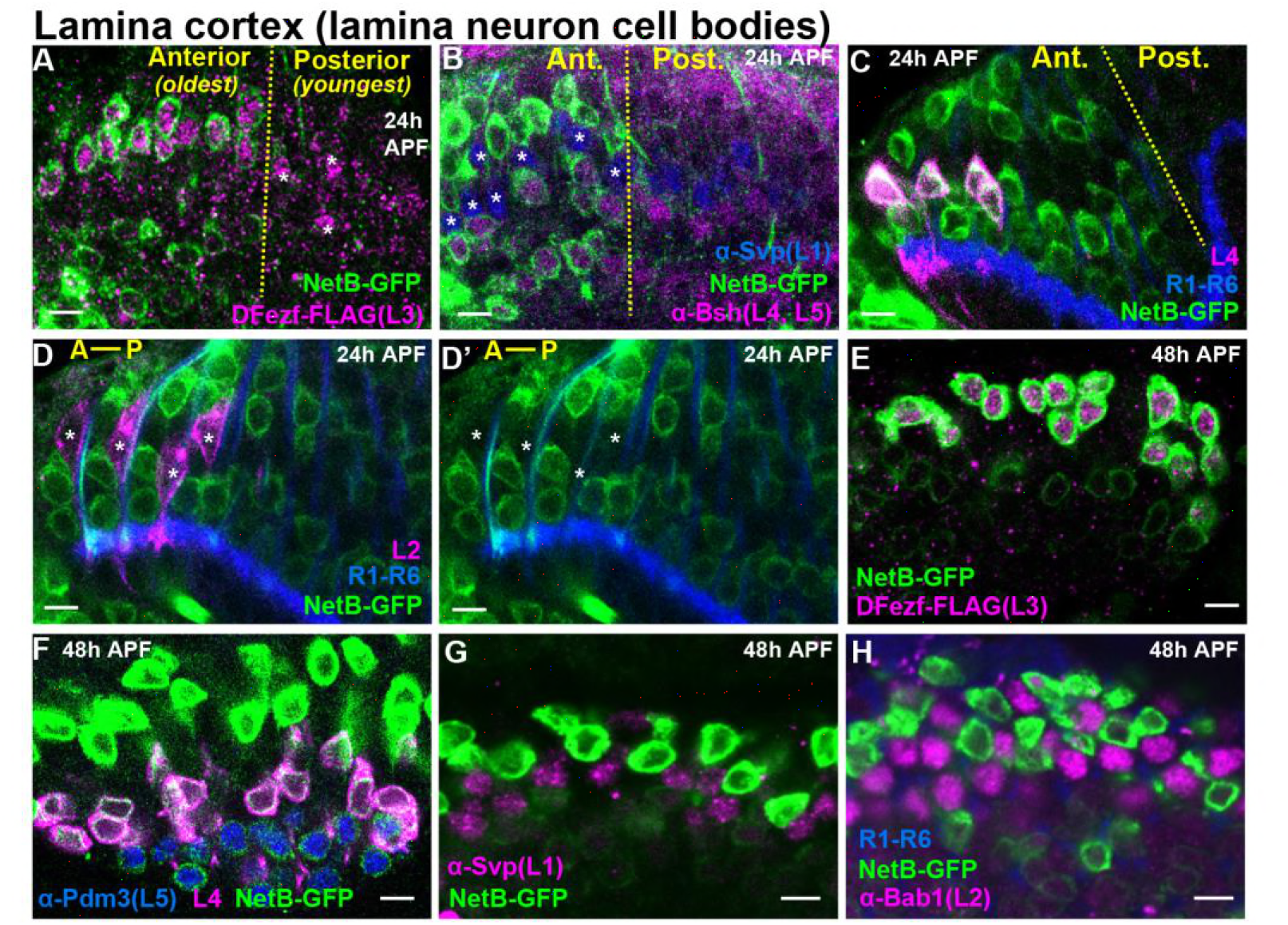
Cellular expression of Net-B in the lamina during pupal development. (A-H) Net-B expression in the lamina at 24 or 48h APF. Endogenous Net-B was visualized using a Net-B-GFP protein trap (α-GFP). All confocal images are representative examples, and at least five brains were examined per genotype. All scale bars = 5 microns (A-D’) In the lamina, lamina neurons are born in a wave (Anterior = oldest, Posterior = youngest). The yellow dashed lines in (A-C) indicate the anterior-posterior region of the lamina where Net-B expression is first observed. In D and D’ anterior-posterior is indicated by A-P. (A) At 24h APF, Net-B (green) was expressed in the anterior side of the developing lamina (to the left of the yellow dashed line) containing the oldest lamina neurons. Net-B was expressed in L3 neurons (magenta, dFezf-FLAG). The youngest L3 neurons on the posterior side of the lamina that had not turned on Net-B expression are indicated by asterisks. (B) At 24h APF, Net-B (green) was detected in L4 and L5 neurons (magenta, α-bsh) in the oldest region of the lamina (to the left of the yellow dashed line). Net-B (green) was not detected in L1 neurons (blue, α-svp, asterisks in the anterior region). (C) Consistent with findings shown in B, at 24h APF Net-B (green) was expressed in the oldest L4 neurons (magenta) in the anterior region of the lamina (left of the dashed yellow line). L4 neurons were labeled using an L4-specific driver (31C06-LexA, LexAop-myr-tdTOM). At this stage 31C06-LexA drives expression in a small population of L4 neurons in the anterior region of the lamina. R1-R6 axons (blue. mAB24B10) provided a reference for the lamina. The image shown here is taken from a pupa that is slightly older than in A and B, as Net-B is expressed in nearly all the lamina neurons in the field of view. (D and D’) Net-B (green) was not expressed in L2 neurons (magenta) at 24h APF. L2 neurons were visualized using a specific driver (16H03-LexA, LexAop-myr-tdTOM) that drives expression in the most anterior (oldest) L2 neurons at this stage. R1-R6 axons (blue, mAB24B10) serve as a reference for the lamina. Asterisks show the positions of L2 somas. The image shown here is taken from a pupa that is slightly older than in A and B, as Net-B is expressed in all the lamina neurons in the field of view. (E) Net-B (green) is expressed in all L3 neurons (magenta, dFezf-FLAG) at 48h APF. (F) All L4 (magenta, 31C06-LexA, LexAop-myr-tdTOM) and L5 (blue, α-Pdm3) neurons express Net-B at 48h APF. Net-B expression in L4 and L5 neurons is weaker than in L3 neurons (green only). (G) Net-B (green) is not expressed in L1 neurons (magenta, α-Svp) at 48h APF. (H) Net-B-(green) is not expressed in L2 neurons (magenta, α-Bab1) at 48h APF.

To test whether dFezf is required for Netrin protein expression we examined Net-B expression during pupal development when dFezf function was disrupted in L3 neurons. Disrupting *dFezf* in single L3 neurons in MARCM experiments drastically reduced Net-B immunostaining in L3 cell bodies at 48h APF (Fig. 6A-B’). To assess whether *dFezf* is necessary for Netrin expression in the M3 layer, wherein it is thought to be secreted from L3 growth cones, we performed RNAi experiments to disrupt *dFezf* in the vast majority of L3 neurons. The expression of *dFezf* RNAi in the lamina eliminated detectable dFezf immunoreactivity in L3 neurons (Fig. 6C and D), and considerably reduced Net-B immunostaining in L3 somas and the developing M3 layer at 48h APF (Fig. 6E and F). Together, these experiments demonstrate that *dFezf* is cell autonomously required for Netrin expression in L3 neurons.

**Figure 6.**
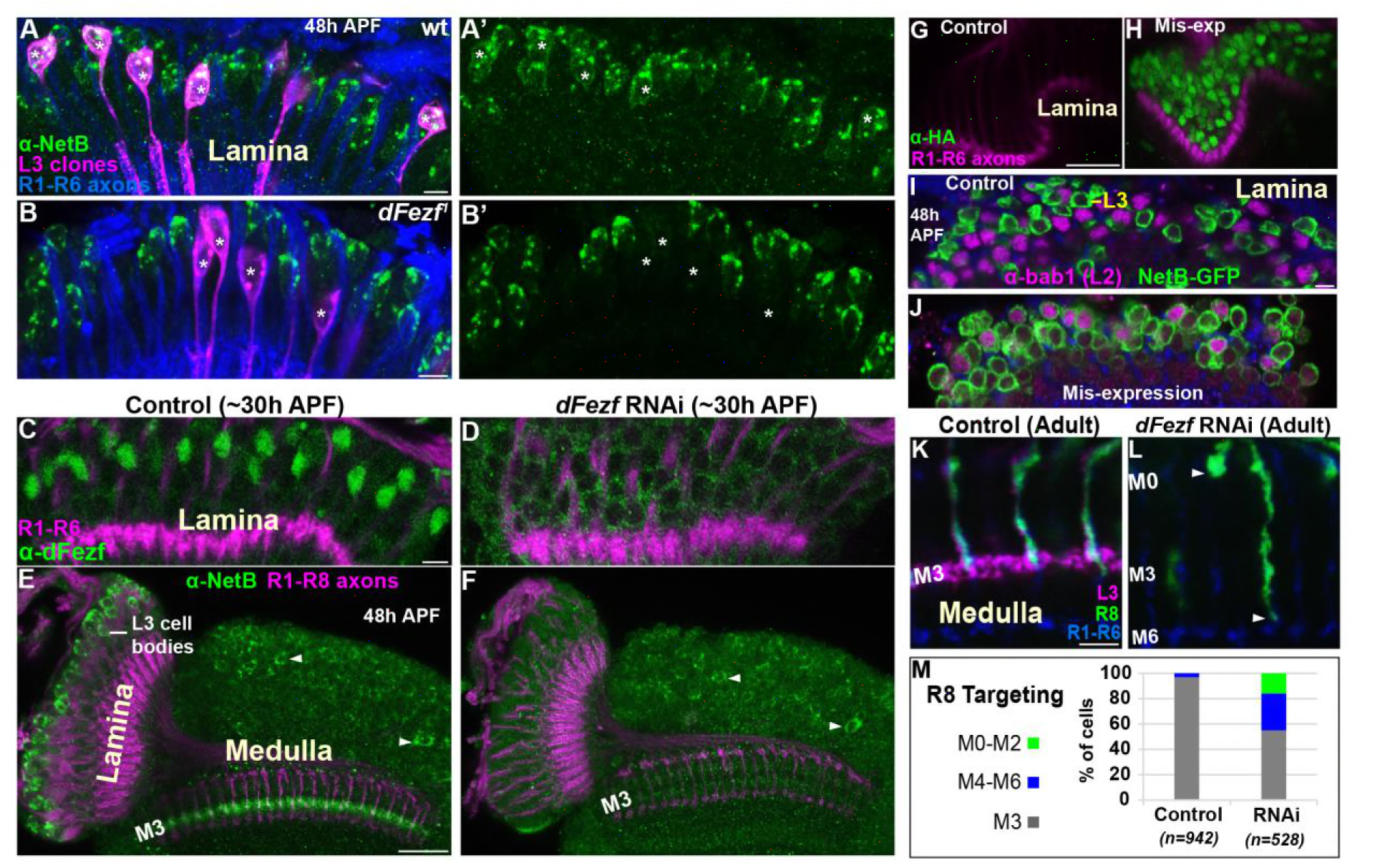
DFezf non-autonomously regulates R8 layer specificity through activation of Netrin expression in L3 neurons. (A-B’) Net-B expression (green, α-Net-B) in the somas of wild type or *dFezf^1^* L3 neurons was assessed at 48h APF in the lamina in MARCM experiments using confocal microscopy. L3 clones (magenta) expressed myr-GFP (9-9-GAL4) and were visualized using a GFP antibody. (A-A’) Asterisks indicate the positions of wild type L3 clones (magenta). The images are representative of Net-B expression (green) assessed in five brains. (A) R1-R6 axons (blue, mAB24B10) were used as a reference for the lamina. Scale bar = 5 microns. (B-B’) Asterisks indicate the positions of *dFezf^1^* L3 clones (magenta). The images are representative of Net-B expression (green) assessed in six brains. (B) R1-R6 axons (blue, mAB24B10) were used as a reference for the lamina. Scale bar = 5 microns. (C and D) *DFezf* RNAi was expressed in lamina neurons and their precursor cells using 9B08-GAL4. DFezf expression (green) in L3 neurons was assessed through immunostaining (α- dFezf). R1-R6 axons (magenta, mAB24B10) provided a reference for the lamina. (C) Representative confocal image of dFezf expression in control flies, which include flies containing *dFezf* RNAi only or 9B08-GAL4 (7 brains total). Scale bar = 5 microns. (D) Shows a representative confocal image of dFezf expression in knockdown flies (8 brains examined). (E and F) Net-B (green, α-Net-B) expression in the lamina and within the M3 layer was assessed in control brains and when *dFezf* RNAi was expressed in lamina neurons and their precursors using 9B08-GAL4. R1-R8 axons (magenta, mAB24B10) provided a reference for the lamina and outer medulla. Arrowheads indicate Net-B expression in medulla neurons. (E) Shows a representative confocal image of Net-B expression (green) in control flies, which contain *dFezf* RNAi only or 9B08-GAL4 (4 brains total). Net-B was prominently observed in L3 cell bodies in the lamina, within the M3 layer in the medulla and in medulla neuron cell bodies (arrowheads) on a consistent basis. Scale bar = 20 microns. (F) A representative confocal image from flies containing *dFezf* RNAi + 9B08-GAL4 (6 brains). Net-B (green) was consistently reduced in L3 cell bodies in the lamina and the M3 layer, but not in medulla neuron cell bodies (arrowheads). (G-J) DFezf-HA was expressed broadly in lamina neurons by 11-164-GAL4. (G and H) Confocal images of dFezf-HA expression (green, α-HA) in lamina neurons (~24h APF). The lamina was identified by visualizing R1-R6 axons (magenta, mAB24B10). (G) Representative confocal image from flies containing DFezf-HA or 11-164-GAL4 only (at least 5 brains). No HA labeling was detected in the lamina of these flies. (H) Shows a representative confocal image from flies containing DFezf-HA + 11-164-GAL4 (at least 5 brains). Prominent HA staining (green) was always observed in the lamina of these flies at 24h APF. (I and J) Confocal images of Net-B expression (green) (NetB-GFP, α-GFP) in the lamina at 48h APF. L2 neurons (magenta) were labeled using α-bab1, and R1-R6 axons (blue, mAB24B10) provided a reference for the lamina. (I) Representative confocal image from flies containing DFezf-HA or 11-164-GAL4 (4 brains). No L2 neurons (magenta) expressed Net-B (green) (n=308). Net-B-expressing cells in the image are L3 neurons. Scale bar = 5 microns. (J) Representative confocal image from flies containing DFezf-HA + 11-164-GAL4 (4 brains). 99% of L2 neurons (magenta) expressed Net-B (green) (n=310). (K-M) *DFezf* RNAi was expressed in lamina neurons and their precursors using 9B08-GAL4 and R8 axon targeting was assessed within the outer medulla. (K and L) Confocal images showing R8 axon targeting within the outer medulla in adult flies. R8 axons (green) were visualized using an R8-specific marker (Rh6-GFP, α-GFP). L3 axons (magenta) are labeled by 22E09-LexA, LexAop-myr-tdTOM. *DFezf* knockdown inhibits 22E09-LexA activity, and serves as a positive control for the efficacy of *dFezf* RNAi. R7 and R8 axons (blue, mAB24B10) provide a reference for outer medulla layers. (K) A representative image from flies containing *DFezf* RNAi or 9B08-GAL4 only (6 brains total). Scale bar = 5 microns. (L) A representative image from flies containing *DFezf* RNAi + 9B08-GAL4 (4 brains). Arrowheads indicate the depths at which R8 axons terminate. (M) Quantification of R8 axon targeting. N = the number of R8 axons counted.

To determine if dFezf is sufficient to induce Netrin expression, we mis-expressed dFezf-HA in lamina neurons using a pan-lamina GAL4 driver (Fig. 6G and H) and assessed Net-B expression in L2 neurons (Fig. 6I and J). L2 neurons mis-expressing *dFezf* still expressed Bab1,a transcription factor shown to be selectively expressed in L2 neurons in the lamina (Tan et al., 2015), indicating that *dFezf* expression did not cause an obvious change in cell fate. We found that, while no wild type L2 neurons expressed Net-B at 48h APF, 99% of dFezf-HA expressing L2 neurons ectopically expressed Net-B. Thus, in the context of L2 neurons dFezf is sufficient to induce Net-B expression. Collectively, our RNA-seq and protein expression experiments support that dFezf, either directly or through an intermediate regulator, activates *Netrin* expression in L3 neurons.

### DFezf in L3 neurons is non-autonomously required for R8 layer specificity

To directly determine if dFezf function in L3 neurons is required for R8 layer specificity, as is predicted by its role in regulating *Netrin* expression, we disrupted *dFezf* via RNAi and assessed R8 axon targeting using an R8-specific marker. Normally, R8 axons terminate in the M3 layer (Fig. 6K, M). When *dFezf* was disrupted 45% of R8 axons terminated within inappropriate layers (Fig. 6L and M). 29% of the axons terminated in layers proximal to M3 (i.e. M4-M6) and 16% terminated in distal layers (i.e. M0-M2). The penetrance of mis-targeting in our experiments is similar to that reported by Timofeev and colleagues in a *Netrin* null background (53%) (Timofeev et al., 2012). However, they reported that only 2% of R8 axons mis-targeted to more proximal layers (versus 29% in our experiments). Differences in R8 mis-targeting between ours and previous experiments could reflect non-specific effects of disrupting dFezf in L3 neurons (e.g. due to mis-targeting L3 axons), or indicate that L3 neurons provide a dFezf-dependent cue in addition to Netrin that is important for the targeting of R8 axons (see discussion).

Nonetheless, our experiments show that *dFezf* in L3 neurons is required for R8 layer specificity. Together with previous research, these findings along with our analyses of Netrin expression indicate that dFezf regulates R8 layer specificity, to a large extent, through activation of *Netrin* expression in L3 neurons.

### Disrupting dFezf in L3 neurons perturbs connectivity with Tm9 neurons

Within the M3 layer L3 axons synapse onto the dendrites of transmedullary 9 neurons (Tm9) (Gao et al., 2008; Takemura et al., 2013; Takemura et al., 2015), and synapses between R8 and Tm9 have also been reported (Gao et al., 2008). Tm9 neurons are a subtype of projection neurons that transmit information from the medulla to the lobula neuropil. As disrupting *dFezf* in L3 neurons caused defects in L3 and R8 layer innervation but did not impair the formation of presynaptic sites in L3 neurons (Fig. 2K and L), we assessed whether connectivity with Tm9 neurons is perturbed under these conditions. To test this we used cell-specific drivers to assess overlap between L3 axons and Tm9 dendrites within the same medulla columns during development, under conditions when L3 neurons were wild type or mutant for *dFezf* in MARCM experiments. During early pupal development (~24h APF) in wild type columns, L3 growth cones and Tm9 dendrites targeted to similar depths within the outer medulla, but appeared to be restricted to opposite sides of the column (Fig. 7A, B-B”). In contrast, by mid-pupal development (~48h APF) L3 growth cones and Tm9 dendrites overlapped significantly within the developing M3 layer (Fig. 7C, D-D”). In adult flies L3 terminals were flattened and overlapped with the proximal portion of Tm9 dendrites within the M3 layer (Fig. 7E, F-F”).

**Figure 7.**
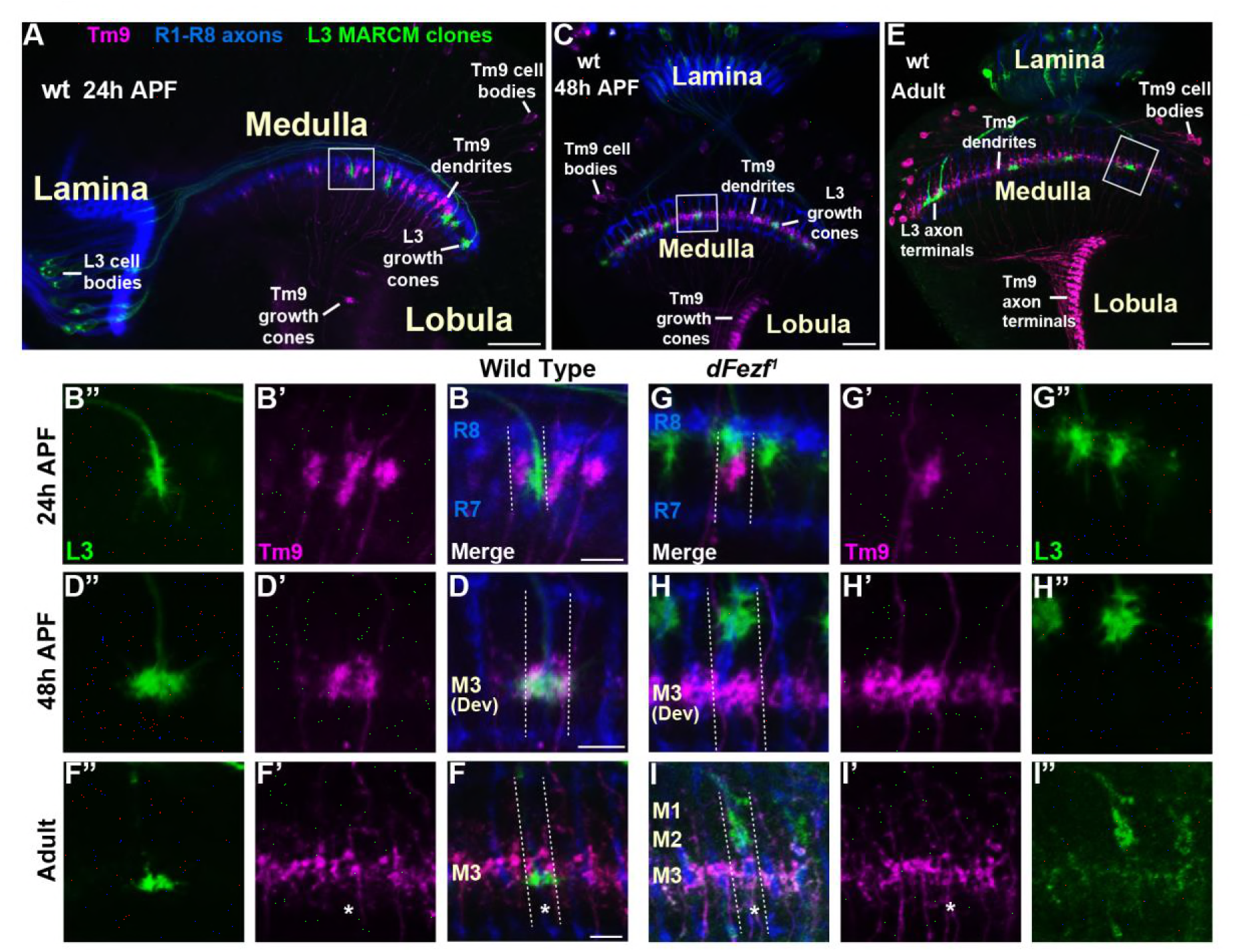
*Disrupting dFezf in L3 neurons perturbs connectivity with Tm9 neurons*. (A-I”) Confocal images from MARCM experiments analyzed at 24h APF, 48h APF or in newly eclosed adults. The images shown are representative of neurons assessed from at least five brains per genotype per time-point. L3 clones (green) expressed myr-GFP (9-9-GAL4) and were visualized using a GFP antibody, wild type Tm9 neurons (magenta) expressed myr-tdTOM (24C08-LexA) and were visualized with a DsRed antibody. R1-R8 axons (blue, mAB24B10) define the lamina and outer medulla. (A, C, E) Broad views showing the entire lamina and medulla and the morphologies of wild type L3 (green) and Tm9 neurons (magenta). The solid boxed regions show examples of the regions of the outer medulla shown in B-B”, D-D”, F-F”, G-G”, H-H”, and I-I”. (B, D, F, G, H, I) Dashed lines indicate the estimated boundaries of a medulla column using R7 and R8 axons (blue) as a reference. (F and F’’, I and I’) Asterisks mark the positions of medulla columns containing L3 clones.

When *dFezf* was disrupted in L3 neurons the positions of L3 growth cones and Tm9 dendrites within the same columns were altered in early pupal development (Fig. 7G-G”). Frequently, the growth cones and dendrites were positioned at different depths within the outer medulla. At mid-pupal development L3 growth cones and Tm9 dendrites did not overlap within the column, with most of the growth cones innervating developing layers in the distal medulla (M1-M2) while Tm9 dendrites innervated the developing M3 layer normally (Fig. 7H-H”). In adult flies, we found that compared to wild type L3 clones, *dFezf^1^* clones were weakly labeled by an L3-specific GAL4 driver (9-9-GAL4) (Fig. 7I, I”). Despite the weak labeling, we observed that L3 axon terminals and Tm9 dendrites innervated different depths within the neuropil, and the morphologies of Tm9 dendrites in columns containing wild type or *dFezf^1^* L3 neurons were indistinguishable (Fig. 7I-I”). The decreased labeling observed in mutant clones is likely due to a reduction in 9-9-GAL4 activity during pupal development rather than cell death, as *dFezf^1^* L3 neurons in adult flies were strongly labeled with a pan-neuronal driver (i.e. the *brp* locus) (see Fig. 2G-I, L, N). Thus, dFezf may be required to maintain 9-9-GAL4 expression in L3 neurons.

To summarize, in these experiments we did not observe overlap between the axons of *dFezf* null L3 neurons and Tm9 dendrites during development or in adult flies, strongly pointing to disrupted connectivity between these neurons under these conditions. In addition, it is also likely that dFezf is partially required for R8-Tm9 connectivity, as a significant number of R8 axons terminate in layers distal to M3 in the absence of dFezf function (Fig. 6L and M), and R8 axons and Tm9 dendrites are unlikely to overlap in these columns.

## DISCUSSION

Laminar organization of synaptic connections is a principal feature of nervous system structure. Determining how cells arrange their connections into layered networks is thus vital for understanding how precise neural connectivity is established. Molecules necessary for layer innervation in isolated contexts have been identified. However, whether overarching mechanisms coordinate cells to the correct layers is unclear. Here we address this by illuminating a transcriptional mechanism that coordinates different cell types to the same layer in the *Drosophila* visual system (summarized in Fig. 8). Based on our findings, we propose that the precision of layer-specific connectivity is controlled by transcriptional modules that cell-intrinsically instruct targeting to specific layers, and cell-extrinsically recruit specific neuron types.

**Figure 8.**
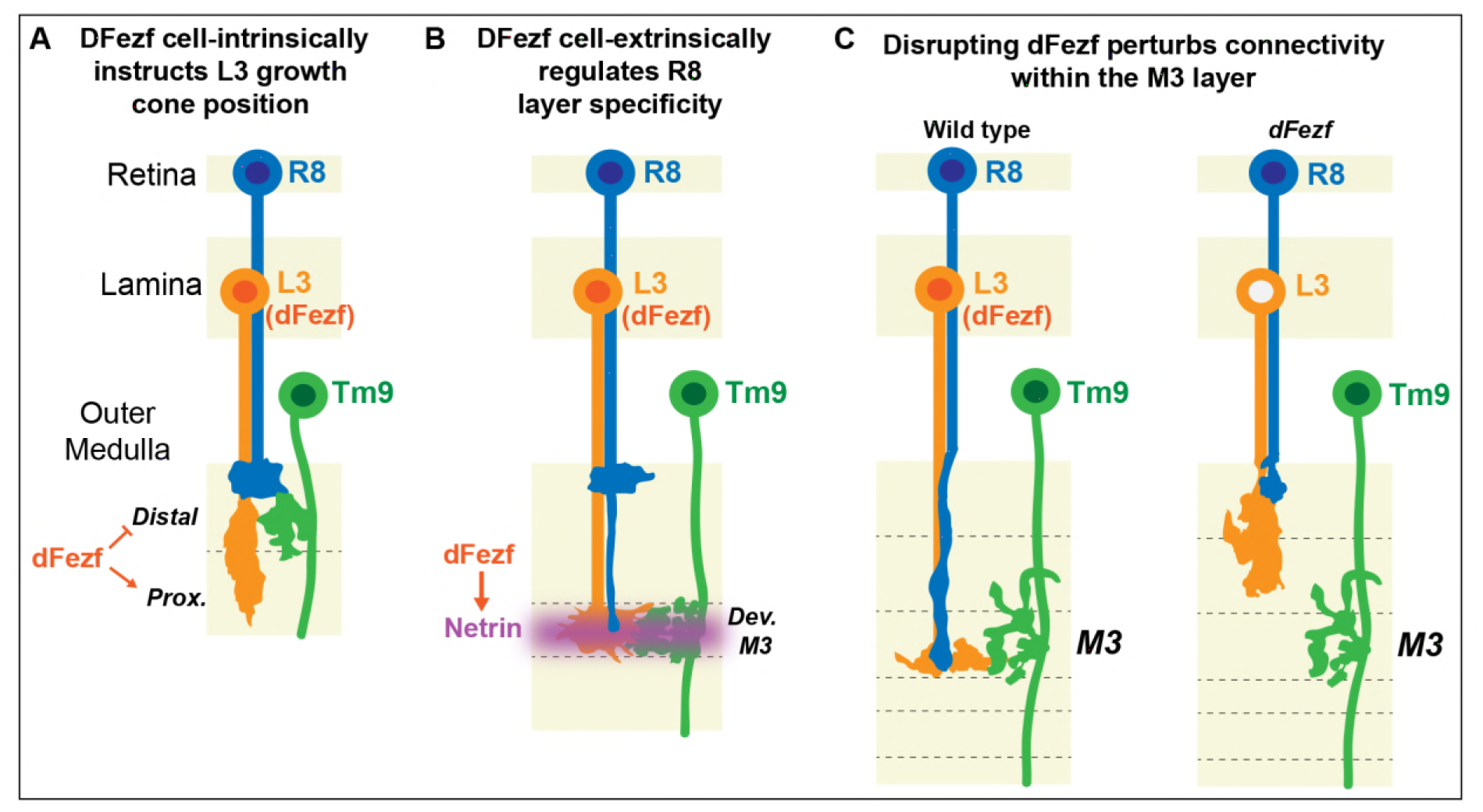
DFezf coordinates the targeting of L3 and R8 axons during development. (A) Early in medulla development, dFezf promotes the targeting of L3 growth cones to the proximal versus distal domain of the outer medulla. (B) L3 growth cones segregate into the developing M3 layer and secrete Netrin, which regulates the attachment of R8 growth cones within the layer. DFezf also regulates this step by activating the expression of *Netrin* in L3 neurons. (C) Within the M3 layer, L3 and R8 axons synapse onto Tm9 dendrites. When dFezf function is lost in L3 neurons, L3 and R8 axons innervate inappropriate layers while Tm9 dendrites innervate the M3 layer normally. As a result, connectivity with Tm9 neurons is disrupted.

### DFezf cell-intrinsically instructs axon target specificity

Before innervating discrete layers, lamina neuron axons terminate within broad domains in the outer medulla (Fig. 1C). The targets of lamina neuron axons within broad domains are unknown, and could include synaptic partners, intermediate targets, guidepost cells, and molecules embedded in the extracellular matrix. Our findings demonstrate that dFezf regulates broad domain specificity in L3 neurons, and suggest that targeting to the correct domain is necessary for proper layer specificity and synaptic connectivity.

*DFezf* function promotes targeting to the proximal domain of the outer medulla, implying that dFezf regulates a program of gene expression that instructs growth cone targeting to this region. DFezf-mediated gene expression could inhibit the recognition of signals in the distal domain of the outer medulla, promote recognition of signals in the proximal domain, or both. In a simple model, dFezf could repress or activate the expression of cell surface receptors that bind ligands in the distal domain or proximal domain, respectively. Receptors that are down regulated by dFezf could normally function to promote the targeting of L2 or L4 growth cones to the distal domain of the outer medulla, while up regulated receptors could also act in L1 and L5 neurons to promote targeting to the proximal domain (although their expression would be activated by alternative factors).

Genetic and expression analyses show that dFezf acts in parallel to CadN and Sema-1a in L3 neurons to regulate growth cone targeting. We envision two models by which these molecules could act in combination in this context. In one model, precise targeting is achieved by preventing innervation of inappropriate regions. Here, dFezf prevents growth cone termination within the distal domain of the outer medulla, while CadN and Sema-1a prevent innervation of the serpentine layer and inner medulla. Thus, L3 growth cones target to the proximal domain of the outer medulla through mechanisms that prevent both superficial and deeper innervation, rather than through a direct targeting mechanism. In the second model, L3 growth cones are directly targeted to the proximal domain of the outer medulla. In this case, dFezf regulates genes that direct targeting to the proximal domain (dFezf could in parallel regulate genes that prevent termination in the distal domain), and CadN and Sema-1a play a role in consolidating growth cone position. Identification and characterization of dFezf target genes in L3 neurons will shed light on the mechanism by which dFezf regulates growth cone targeting. Moreover, as dFezf is expressed in L3 neurons throughout development, such studies could provide insight into the mechanisms underlying subsequent steps in layer connectivity, such as segregation into the developing M3 layer and synaptic specificity within the layer.

### DFezf regulates layer innervation through a cell-extrinsic mechanism

In the absence of *dFezf* function in L3 we observed R8 axons inappropriately innervating distal layers (i.e. M0-M2), as expected in the absence of Netrin function (Akin and Zipursky, 2016; Timofeev et al., 2012), and a significant fraction of R8 axons terminating in proximal outer medulla layers (M4-M6), which was unexpected. In vivo time lapse imaging studies showed that the axons of *Frazzled* (Netrin receptor) null R8 neurons targeted to M3 normally and then retracted, indicating that Netrin/Frazzled signaling is required for growth cone stabilization *within* the target layer, but is dispensable for recognition *of* the target layer (Akin and Zipursky, 2016). Thus, additional molecules regulate R8 layer specificity. One interesting possibility is that in addition to *Netrin,* dFezf regulates the expression of a gene(s) important for R8 axons to “recognize” the M3 layer, and in the absence of this cue a subset of R8 axons overshoot M3 and innervate proximal layers. Indeed, when L3 neurons were genetically ablated a significant percentage of R8 axons terminated in proximal outer medulla layers (Pecot et al., 2014), consistent with L3 growth cones providing a “stop signal” to extending R8 growth cones. This idea could be tested in future studies by identifying cell surface and secreted molecules downstream of dFezf in L3 neurons, and assessing whether disrupting these genes in L3 causes defects in R8 layer specificity. In addition, it will be important to assess whether dFezf targets regulate the layer innervation of multiple cell types or specifically regulate R8 layer specificity, and to elucidate the mechanisms by which dFezf regulates effector gene expression (e.g. directly or through intermediate factors).

### A transcriptional mechanism for controlling the stepwise formation of layer-specific circuits

Previous studies indicate that medulla layers are not pre-established regions that serve as a template for circuit formation, but rather that layers are built over time from broad domains as the neurites of different cell types are added in a precise order. Thus, central to elucidating the molecular and cellular logic underlying assembly of the medulla network is identifying the molecular and cellular components involved in the sequence of interactions giving rise to specific layers, and discovering commonalities or connections between interactions and how different layers are assembled. Insight into the cellular mechanisms giving rise to the M3 layer circuitry was provided by the discovery that R8 innervation of M3 relies on a signal from L3 axons (Netrin) (Pecot et al., 2014; Timofeev et al., 2012), which innervate M3 at an earlier stage despite being born later in development (Selleck and Steller, 1991). These studies and the fact that L3 and R8 neurons contribute input to common pathways suggest that the cellular mechanisms governing circuit formation reflect functional relationships between neurons, rather than birth order.

Less progress has been made in identifying the molecular logic underlying the stepwise assembly of layer-specific circuits. This is likely due in part, to the fact that how neurons innervate particular layers is cell type-specific, rather than layer-specific. For example, while R8 neurons depend on Netrin for layer specificity, Netrin is dispensable for the layer innervation of L3 and Tm9 neurons (unpublished results). Thus, identifying commonalities in the mechanisms used by neurons to innervate layers has been challenging. However, here we show that L3 and R8 layer innervation is linked through dFezf, which controls a program of gene expression that regulates the layer innervation of both neurons. Thus, while L3 and R8 utilize different molecules to achieve layer specificity, the expression of key molecules required by each neuron is controlled by the same transcription factor.

Using the same transcriptional pathway or module, to control genes that function cell-intrinsically and cell-extrinsically to regulate layer innervation, represents a simple mechanism for controlling the series of interactions that lead to circuit formation. Employing specific transcriptional modules in specific neurons would ensure that those neurons innervate the correct target layer, and also express the genes necessary for the targeting of other circuit components. We hypothesize that this strategy is widely employed to regulate the assembly of circuitry within the medulla. For example, cell-specific transcription factors analogous to dFezf in L3 have been identified in each of the other lamina neuron subtypes (Tan et al., 2015), and the positions of lamina neuron arborizations in the medulla are well suited to produce signals that regulate the precise layer innervation of other cell types (Fig. 1C). Indeed, CadN-based interactions between L2 terminals and L5 axons are necessary for L5 branching within the M2 layer (Nern et al., 2008). Thus, we hypothesize that the function of cell-specific transcriptional modules in lamina neurons coordinates the assembly of discrete layer-specific circuits within the outer medulla.

We also speculate that the function of dFezf in L3 neurons is conserved in laminated regions of the mammalian nervous system. For instance, in the inner plexiform layer (IPL) of the mouse retina, which is analogous to the medulla in structure and function (Sanes and Zipursky, 2010), Fezf1 is expressed in a subset of bipolar cells that innervate specific sublaminae (i.e. layers) (Shekhar et al., 2016). Bipolar cells are analogous to lamina neurons, and we hypothesize that Fezf1 in specific types of bipolar cells functions analogously to dFezf in L3 to coordinate laminar-specific connectivity.

In the cerebral cortex of mice, Fezf2 cell-intrinsically controls the identity of subcortically projecting pyramidal neurons that predominantly reside in layer V (Chen et al., 2005a; Chen et al., 2005b; Molyneaux et al., 2005), and pyramidal neuron identity was shown to be important for the laminar positioning of specific classes of GABAergic neurons (Lodato et al., 2011). While it remains unclear as to whether Fezf2 regulates the expression of molecules that recruit specific inhibitory neurons, one interesting possibility is that Fezf2 in the cortex organizes laminar-specific circuitry through cell-intrinsic and cell-extrinsic mechanisms similar to dFezf in the medulla. In conclusion, we expect our findings reflect an evolutionarily shared strategy for assembling laminar-specific neural circuits across species.

## ACKNOWLEDGEMENTS

This research has been aided by grants from the Lefler Center for the Study of Neurodegenerative Disorders (M.Y.P, J.P.) and the Howard Hughes Medical Institute (Gilliam Fellowship for Advanced Study, I.J.S.). We acknowledge C.Y. Lee and H. Wang for providing transgenic flies and the dFezf antibody, respectively. We would also like to thank Victor Barrera of the Harvard Chan Bioinformatics Core, Harvard T.H. Chan School of Public Health, Boston, MA for assistance with the RNA-seq analysis. VB’s work was funded by the Harvard NeuroDiscovery Center. Finally, we acknowledge D.D. Ginty, Y. Chen, G. Fishell, C.C. Harwell and C.L. Cepko for valuable discussions.

## AUTHOR CONTRIBUTIONS

M.Y.P., I.J.S., J.P. and M.S. designed research. I.J.S. performed MARCM, mis-expression, developmental expression, and LN marker expression experiments, and contributed to RNA seq experiments. J.P. executed MARCM, BAC rescue, Netrin, and RNA seq experiments. C.A. and A.K. performed STaR and MARCM experiments. B.G. and K.S. performed experiments assessing L3-Tm9 connectivity. C.X. contributed valuable input to the manuscript. M.Y.P. contributed to MARCM, Netrin, and RNA seq experiments, directed the research and wrote the manuscript with input from the other authors.

## MATERIALS AND METHODS

A list of the complete experimental genotypes in each figure can be found in Table S1 pg. 50-55).

### Experimental model and subject details

#### Description of replicates

In general, the replicates for each experiment are indicated by the number of cells and brains examined (denoted in the figures and figure legends). There are essentially three types of replicates we consider. Within each optic lobe, each column is replicated hundreds of times, different lobes of the same brain are replicates, and the lobes of different brains are also replicates. In all experiments, the phenotypes reported are consistently reproducible for each type of replicate considered (i.e. within and between different animals)

#### Fly strains

Flies were raised on standard cornmeal-agar based medium. Male and females flies were used at the following development stages: 3^rd^ instar, 12 hour after pupariam formation (h APF), 24h APF, 48h APF, 72h APF and newly eclosed adults (within 5 hours of eclosion).

The following strains were obtained from Bloomington Stock Center (Indiana University):

10xUAS-IVS-myr::GFP [attp2], tubP-GAL80, FRT40A, R27G05-FLPG5.PEST [attP40], 20XUAS-RSR.PEST [attP2], 10XUAS-IVS-myr::GFP [su(Hw)attP8], GMR16H03-lexA [attP40], 10xUAS-FRT-stop-FRT-myr::GFP [su(Hw)attP5], UAS-Dcr-2, GMR24C08-lexA [attP40], GMR9B08-GAL4 [attP2], TRiP.JF02342 [attP2] (dFezf RNAi), 20xUAS-RSR.PEST [attP2], Rh6-EGFP

The following strains were obtained from Janelia Research Campus:

27G05-FLP (X), 22E09-LexA

The following stocks were generated in the Pecot Laboratory for this study:

18B02-dFezf, 18B02-dFezf-C1-3xFLAG

The following strains were gifts from other laboratories:

9-9-GAL4, 6-60-GAL4, 11-164-GAL4 (U. Heberlein), LexAop-myr::tdTomato [su(Hw)attP5] (S.L. Zipursky), UAS-dFezf-3xHA, dFezf^1^, dFezf^2^ (C.Y. Lee), CadN1-2 ^Δ14^ (T.R. Clandinin)

### Construction of transgenic animals

#### Generation of the BRP presynaptic marker 79C23S-RSRT-STOP-RSRT-smFPV5-2A-LexA

Restriction free cloning (van den Ent and Löwe 2006) was used to insert smFPV5 (Viswanathan et al., 2015) (addgene: pCAG_smFPV5 plasmid# 59758) in between RSR and the 2A peptide (2A) within “cassette F” (see below), which was made by Chen and colleagues for generation of the original Brp STaR marker (Chen et al., 2014). This replaced the original V5 tag and generated:

Cassette G: GS linker-RSR-STOP-RSR-smFPV5-2A-LexAVP16 [Cassette F: GS linker-RSR-STOP-RSR-V5-2A-LexAVP16]

The modification of the BAC CH321-79C23S (Chen et al., 2014) was performed using the recombineering protocol described previously (Sharan et al., 2009, Nature Protocols).

#### Generation of 18B02 dFezf-C1-3xFLAG

Bacterial artificial chromosome (BAC) with endogenous dFezf locus CH321-18B02 was ordered from BACPAC Resources. The modification of the BAC was performed using the recombineering protocol described previously (Sharan et al., 2009, Nature Protocols).

The wild type 18B02 BAC was transformed into the SW102 E. coli strain (Warming et al., 2015, NAR). RpsL-Kan cassette with homology arms flanking the first stop codon of the dFezf locus was amplified using the following primers:

Forward:

ACCATCAGCAGCAGCAAAGACTCTCGGAGACCTTCATAGCCAAGGTGTTTAGCTTCACGCTGCCGCAAGCACTCAG

Reverse:

GAGGTCGACCCCGTGGAGCTGTTCAAGCTTTCCGCATAGTATCGCTGTCAGGGGTGGGCGAAGAACTCCAGCATGA

The PCR product was transformed into the 18B02-harboring SW102 cells, and the RpsL-Kan cassette was inserted before the first stop codon by homologous recombination.

1kb of DNA fragment centered by the first stop codon of the dFezf locus was PCR amplified from the genomic DNA using the following primers:

Forward: CTCACCTTCCACATGCACAC Reverse: GCGTGTCTACACGGAACTCA

This fragment was then cloned into pGEM-T vector (Promega). Using the resulting construct as template, site-directed mutagenesis was performed by PCR using the following primers so that the 3xFLAG sequence was inserted before the first stop codon of the dFezf locus on the plasmid:

Forward:

GAGACCTTCATAGCCAAGGTGTTTGACTACAAAGACCATGACGGTGATTATAAAGATCATGACATCGATTACAAGGATGACGATGACAAGTGACAGCGATACTATGCGGAAAGC

Reverse: CGAGAGTCTTTGCTGCTGCTGATG

Using the resulting construct as the template, PCR was performed using the following primers to amplify the DNA fragment having 3xFLAG inserted before the first stop codon of the dFezf locus and flanked by ~500 bp of homology arms:

Forward: CATCCGCAAAGCCAGCAACA

Reverse: GAGGTGGCCATCGAGGATAT

The PCR product was then transformed into the SW102 cells that harbored modified 18B02 with RpsL-Kan cassette inserted before the first stop codon of the dFezf locus. The RpsL-Kan cassette was then replaced by 3xFLAG by recombination, resulting in a BAC that has 3xFLAG inserted right before the first stop codon of the dFezf locus. The 3xFLAG tagged BAC was inserted into the VK33 genomic site.

## METHOD DETAILS

### Immunohistochemistry

Fly brains were dissected in Schneider’s medium and fixed in 4% paraformaldehyde in phosphate buffered lysine for 25 minutes. After fixation, brains were quickly washed with phosphate buffer saline (PBS) with 0.5% Triton-X-100 (PBT) and incubated in PBT for at least 2 hours at room temperature. Next, brains were incubated in blocking buffer (10% NGS, 0.5% Triton-X-100 in PBS) overnight in 4 degrees. Brains were then incubated in primary antibody (diluted in blocking buffer) at 4 degrees for at least two nights. Following primary antibody incubation, brains were washed with PBT three times, 1 hour per wash. Next, brains were incubated in secondary antibody (diluted in blocking buffer) at 4 degrees for at least two nights. Following secondary antibody incubation, brains were washed with PBT two times, followed by one wash in PBS, 1 hour per wash. Finally, brains were mounted in SlowFade Gold antifade reagent (Thermo Fisher Scientific, Waltham, MA).

Confocal imaging was accomplished using either a Leica SP8 laser scanning confocal microscope or an Olympus FV1200 Laser Scanning Microscope. Fiji (Schindelin et al., 2012; Schneider et al., 2012) was used to create z stack images.

The primary antibodies used were as follows: anti-GFP (chicken, 1:1000, ab13970) and anti-HA (mouse, 1:1000, ab1424) were purchased from Abcam. Anti-V5 (mouse, 1:200, MCA2892GA) was purchased from Bio-Rad. Anti-HA (rabbit, 1:1000, 3724S) was purchased from Cell Signaling Technologies. Anti-DsRed (rabbit, 1:200, 632496) was purchased from Clontech Laboratories, Inc. Anti-chaoptin (mouse, 1:20, 24B10), anti-elav (rat, 1:200, 7E8A10), anti-CadN (rat, 1:20, DN-Ex 8) and anti-seven-up (mouse, 1:10, 2D3) were purchased from Developmental Studies Hybridoma Bank. Anti-FLAG (mouse, 1:1000, F1804) was purchased from Sigma-Aldrich. Anti-dFezf (rabbit, 1:100) was a gift from C.Y. Lee. Anti-bsh (guinea pig, 1:200) was a gift from S.L. Zipursky. Anti-pdm3 (guinea pig, 1:500) was a gift from J.R. Carlson. Anti-bab1 (rabbit, 1:200) was a gift from S.B. Carroll. Anti-NetB (guinea pig, 1:50) was a gift from B. Altenhein.

The secondary antibodies used were as follows: Goat anti-chicken IgG (H+L) Alexa Fluor 488 (A-11039), Goat anti-rabbit IgG (H+L) Alexa Fluor 568 (A-11011), Goat anti-mouse IgG (H+L) Alexa Fluor 647 (A-21236), Goat anti-mouse IgG1 Alexa Fluor 568 (A-21124), Goat anti-guinea pig IgG (H+L) Alexa Fluor 568 (A-11075), Goat anti-Rat IgG (H+L) Alexa Fluor 647 (A-21247) were all purchased from Thermo Fischer Scientific and were all used at a dilution of 1:500.

### MARCM and MARCM + STaR experiments

In all MARCM experiments, Flp recombinase was expressed in lamina neuron precursor cells using R27G05-FLPG5.PEST (attP40) to induce mitotic recombination, generating single lamina neuron clones, which were visualized using cell-specific GAL4 drivers.

In experiments utilizing STaR and MARCM Flp recombinase in lamina neuron precursor cells induced mitotic recombination generating a subset of L3 neurons that were homozygous for the control chromosome (FRT40) or *dFezf*^1^. In L3 clones R recombinase was expressed (9-9-GAL4) and induced recombination within the Brp locus in a bacterial artificial chromosome (79C23S) resulting in the incorporation of smFPV5 at the C-terminus. In addition, this resulted in the cotranslation of LexA, which activated the expression of LexAop-myrTdTOM to visualize the morphology of L3 clones. Together, this allowed visualization of L3 morphology and endogenous pre-synaptic sites with single cell resolution in wild type conditions or when *dFezf* was disrupted, in an otherwise normal fly.

smFP10V5 contains ten V5 epitopes built into a superfolding GFP backbone (Viswanathan et al., 2015). We find that it is considerably more sensitive than the original V5 tag in the STaR Brp constructs.

### FlpOut experiments

To generate sparsely labeled L5 neurons, we used an interruption cassette (UAS-FRT-STOP-FRT-myrGFP) to drive stochastic expression of myrGFP with an L5 GAL4 driver (6-60-GAL4). Stochastic expression was accomplished through inefficient expression of Flp recombinase in lamina neurons (27G05 X). For reasons that are unknown, this Flp source results in stochastic recombination allowing for visualization of single neurons.

### RNA Seq experiments

For each genotype (control, *dFezf^1^*) 120 crosses were set, with 15 females and 7 males in each cross. After three days, each cross was flipped every day for a week. From those vials we picked out Tb-negative white pre-pupae that formed within a one-hour window and placed the pupae in separate vials. GFP-positive pupae were scored under the fluorescence microscope and removed from the vials. The rest of the pupae were kept at 25°C. After 40h, we dissected the staged pupae within an hour in cold Complete Schneider’s Medium (CSM: 50 mL of CSM consists of 5 mL of heat inactivated fetal bovin serum, 0.1 mL Insulin, 1 mL PenStrep, 5 mL L-Glutamine, 0.4 mL L-Glutathione and 37.85 mL Schneider’s medium, filter setrilized). About 50100 brains were dissected for each independent replicate experiment. After dissection of the brains, the optic lobes were collected and pooled together in 1x Rinaldini’s solution (100 mL Rinaldini’s solution consists of 800 mg NaCl, 20 mg KCl, 5 mg NaH2PO4, 100 mg NaHCO3 and 100 mg Glucose in H2O, filter setrilized) for subsequent dissociation. We used a modified dissociation protocol based on one described previously (Harzer et al., 2013, Nature Protocol). The pooled lobes were washed once with 500 uL 1x Rinaldini’s solution and then incubated at 37°C for 15 min in dissociation solution that consists of 300 uL papain (100 U/mL) and 4.1 uL Liberase at a final concentration of 0.18 Wu/mL. After the enzyme digestion, the lobes were washed twice with 500 uL of 1x Rinaldini’s solution and then twice with 500 uL of CSM. The lobes were then disrupted in 200 uL of CSM by pipetting up and down until the solution became homogeneous. The cell suspention was then filtered through a 30 um mesh 5 mL FACS tube, and diluted with another 500 uL of CSM. The FACS sorting was done on the BD FACS Aria with a nozzle size of 70 um. GFP and dsRed double positive cells were sorted directly into 300 uL of RLT buffer (Qiagen, RNeasy kit) containing 1:100 beta-mercaptoethanol. Total RNA was purified with Qiagen RNeasy kit and eluted in 15 uL of water. After all the samples from 5 replicates (each genotype) were available, we concentrated the RNA into 5 uL by speed vac. 1.5 uL of the concentrated RNA was used to make cDNA library by Smart-seq2 as previously described (Picelli et al., 2014). The quality control of the libraries was done on the Bioanalyzer by Tapestation. The pooled libraries were then loaded onto a single flow cell and sequenced by NextSeq High Output. Between five and fourteen million mapped reads were obtained for each sample.

## QUANTIFICATION AND STATISTICAL ANALYSES

### General

The quantification for immunohistochemistry experiments can be found either within the figure or in the figure legend. In all cases, N refers to the number of cells scored.

### Analysis of RNA seq data

The quality of the sequencing files was initially assessed with the FastQC tool. They were then trimmed to remove contaminant sequences (like polyA tails), adapters and low-quality sequences with cutadapt. These trimmed reads were aligned to the drosophila genome (Flybase relase dmel_r6.15_FB2017_02) using STAR aligner. Alignments were checked using a combination of FastQC, Qualimap, MultiQC and custom tools. Counts of reads aligning to known genes were generated by featureCounts. In parallel, Transcripts Per Million (TPM) measurements per isoform were generated using Salmon. Differential expression at the gene level was called with DESeq2. We used the counts per gene estimated from the Salmon by tximport as input to DESeq2 as quantitating at the isoform level has been shown to produce more accurate results at the gene level. As described in the DESeq2 documentation, the differential expression is assessed using a Wald Test and p-values are adjusted for multiple testing using Benjamini-Hochberg FDR correction.

---

**Figure S1.**
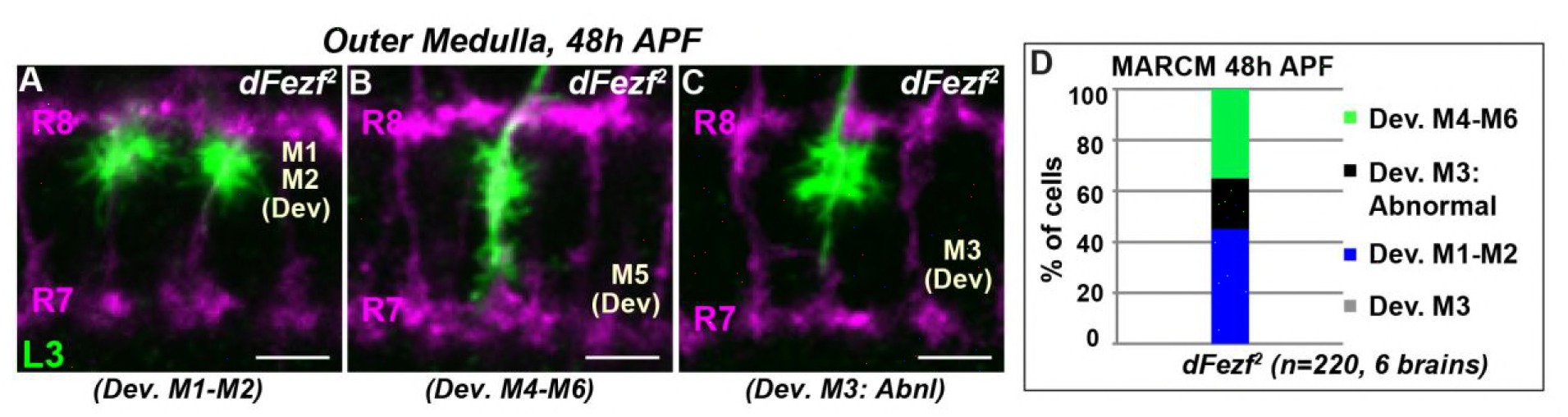
*DFezf^2^* L3 neurons show mis-targeting phenotypes similar to those caused by the *dFezf^1^* allele. (A-C) As in Figure 3, MARCM experiments were performed to generate *dFezf^2^* L3 clones (green) labeled by myr-GFP driven by 9-9-GAL4. At 48h APF, L3 clones are visualized by confocal microscopy using a GFP antibody. Confocal images show L3 growth cones within the outer medulla which is defined by R7 and R8 axons (magenta, mAB24B10). The abnormal targeting and morphologies of the growth cones of mutant neurons resembles that observed for the growth cones of *dFezf^1^* neurons (Figure 3 H-J). All scale bars = 5 microns. (D) Quantification of MARCM experiments at 48h APF. N is the number of neurons that are quantified.

**Figure S2.**
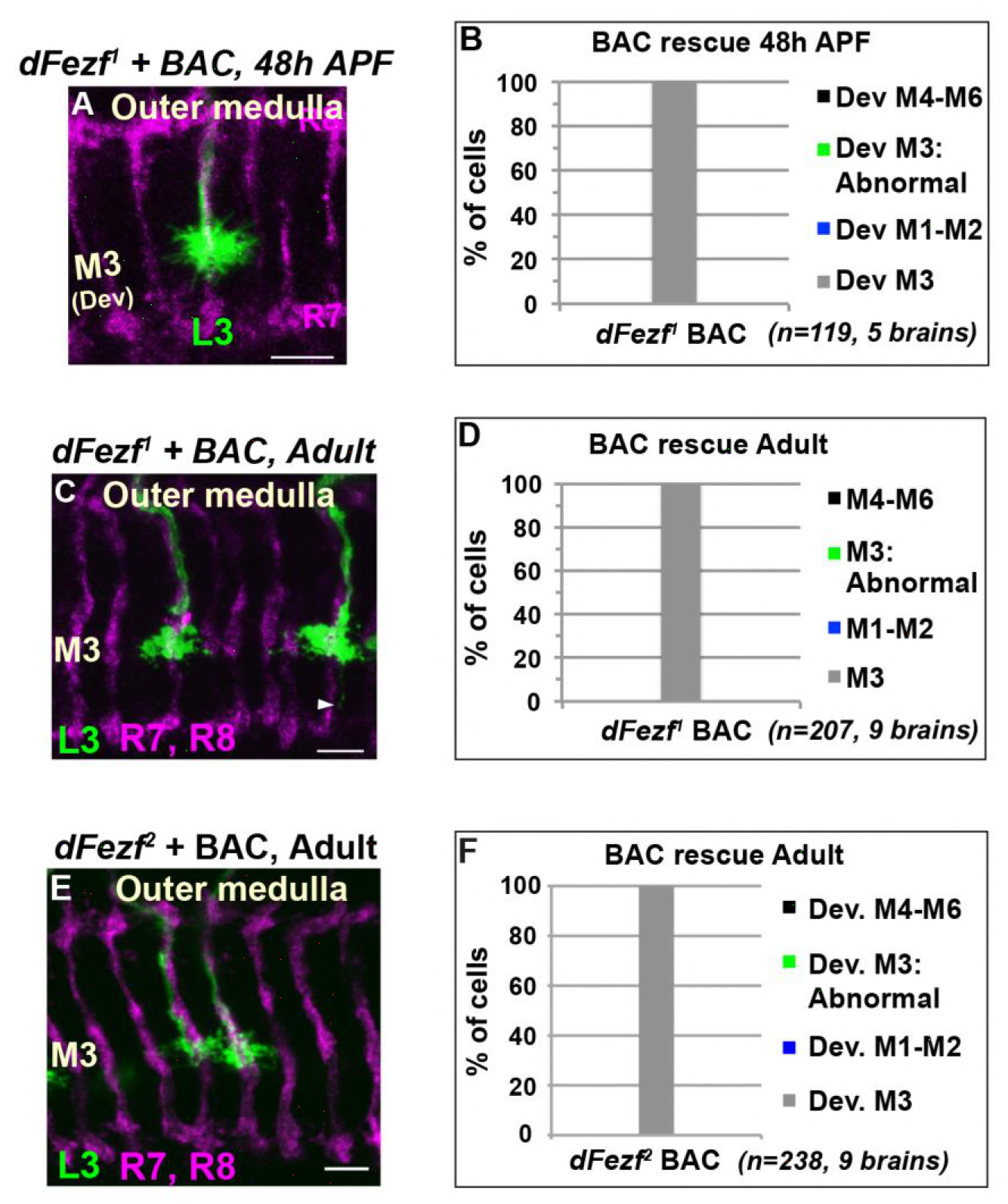
A BAC containing the dFezf locus rescues defects in growth cone targeting, layer specificity and axon terminal morphology caused by *dFezf* null mutations. (A-F) MARCM experiments analyzed at 48h APF or just after eclosion (Adult). L3 neurons are made homozygous for *dFezf^1^* or *dFezf^2^* in the presence of a single copy of a bacterial artificial chromosome (BAC) containing the *dFezf* locus (18B02). (A, C, E) Confocal images of L3 clones (myr-GFP driven by 9-9-GAL4) in the outer medulla. R7 and R8 axons (magenta, mAB24B10) define the boundaries of the outer medulla (24 and 48h APF) or provide a reference for outer layers (adult). Scale bars = 5 microns. (B, D, F) Quantification of MARCM experiments. N = the number of cells counted. (C) Arrowhead indicates a thin process from an L3 terminal extending toward the M6 layer. ~30% of L3 clones in this experiment display this minor abnormality. This likely reflects a partial defect in growth cone retraction from the proximal outer medulla.

**Figure S3.**
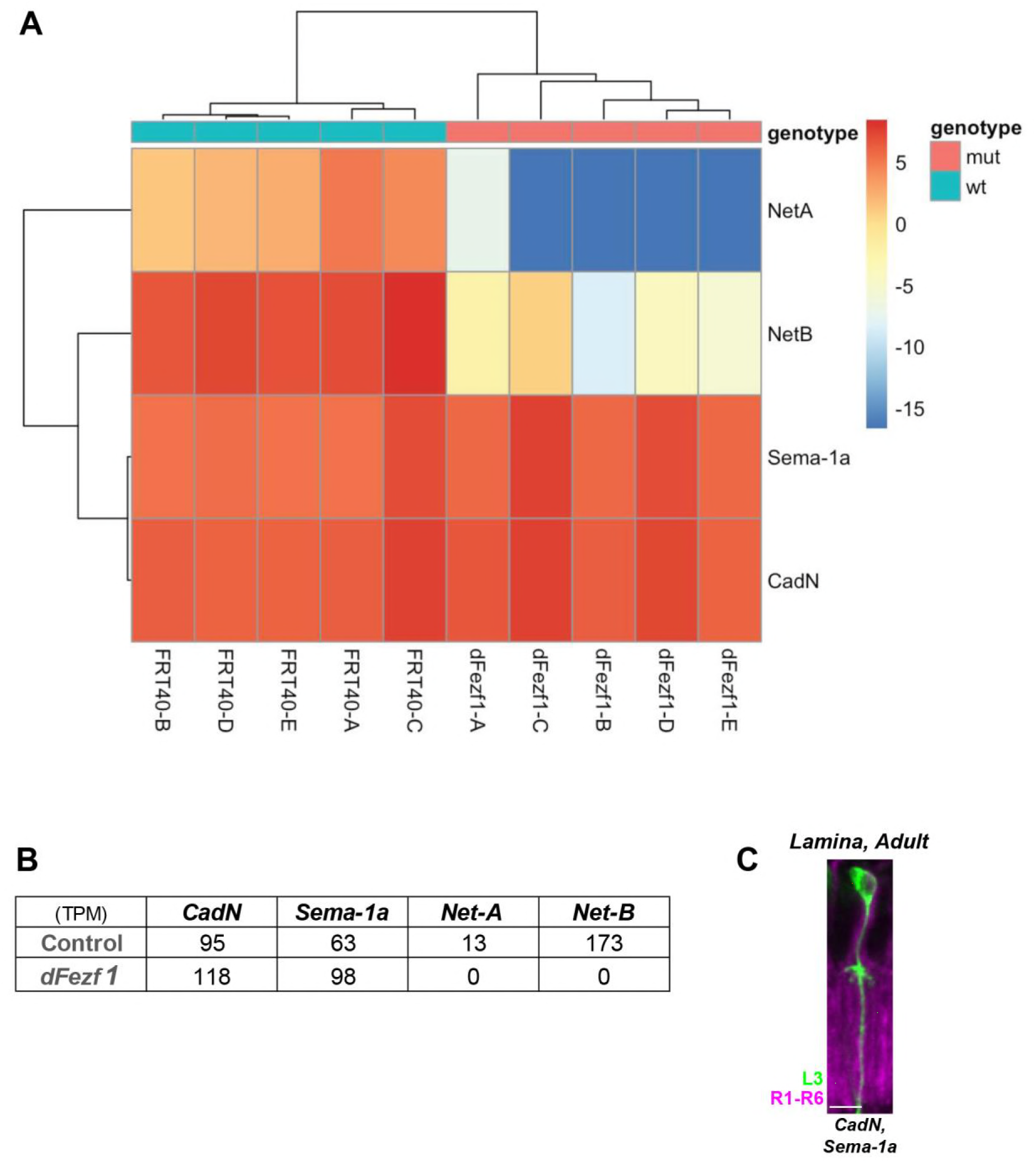
DFezf functions in parallel to CadN and Sema-1a to regulate L3 growth cone targeting. (A and B) The transcriptomes of wild type or *dFezf^1^* L3 clones (generated using MARCM) at 40h APF was determined using RNA-seq (see methods section for a detailed description). Briefly, wild type or *dFezf^1^* L3 clones expressing H2A-GFP (nuclear localization) and myr-tdTOM were dissociated and isolated via FACS. Five biological replicates were performed for each genotype. All libraries were made at the same time, and the samples were sequenced at the same time within the same flow cell (Illumina Next-Seq high output). (A) Heatmap of normalized counts for *CadN*, *Sema-1a*, *Net-A* and *B* after applying an rlog transformation. This is a variance-stabilizing transformation, which transforms the count data to the log2 scale in a way that minimizes differences between samples for rows with small counts, and which normalizes with respect to library size. (B) The average transcript per million (TPM) counts for *CadN*, *Sema-1a*, *Net-A* and *B* are shown for wild type and *dFezf^1^* L3 neurons (across all biological replicates). (C) Disrupting *Sema-1a* and *CadN* in L3 neurons in MARCM experiments (as in Fig. 3Q) drastically reduces L3 dendritic branches. L3 clones expressed myr-GFP driven by 9-9-GAL4 and were visualized in confocal images using a GFP antibody. R1-R6 axons (mab24B10) were used as a reference for the lamina.

**Figure S4.**
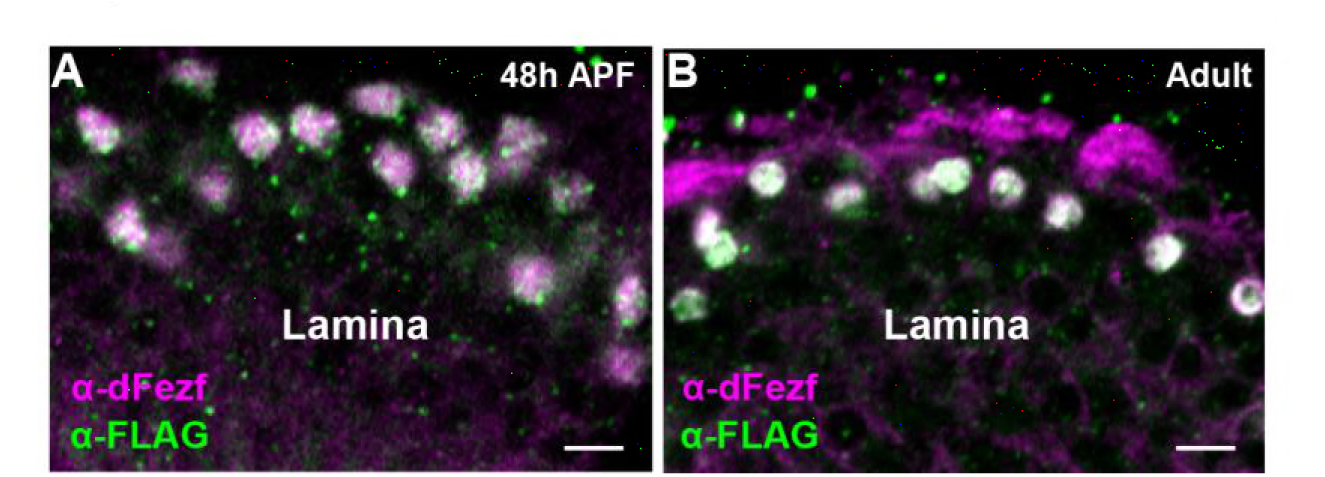
Characterization of dFezf-FLAG (BAC). (A and B) A BAC (18B02) containing the *dFezf* locus was modified to incorporate sequence encoding three consecutive FLAG epitopes (3xFLAG) at the C-terminus prior to the translational stop. DFezf-FLAG (magenta) was detected in L3 somas by confocal microscopy using a FLAG antibody at 48h APF or in newly eclosed flies. All dFezf-FLAG (magenta) expressing cells also expressed endogenous dFezf (green, α-dFezf), and all cells expressing endogenous dFezf expressed dFezf-FLAG. This confirms that dFezf-FLAG recapitulates the normal pattern of dFezf expression in the lamina.

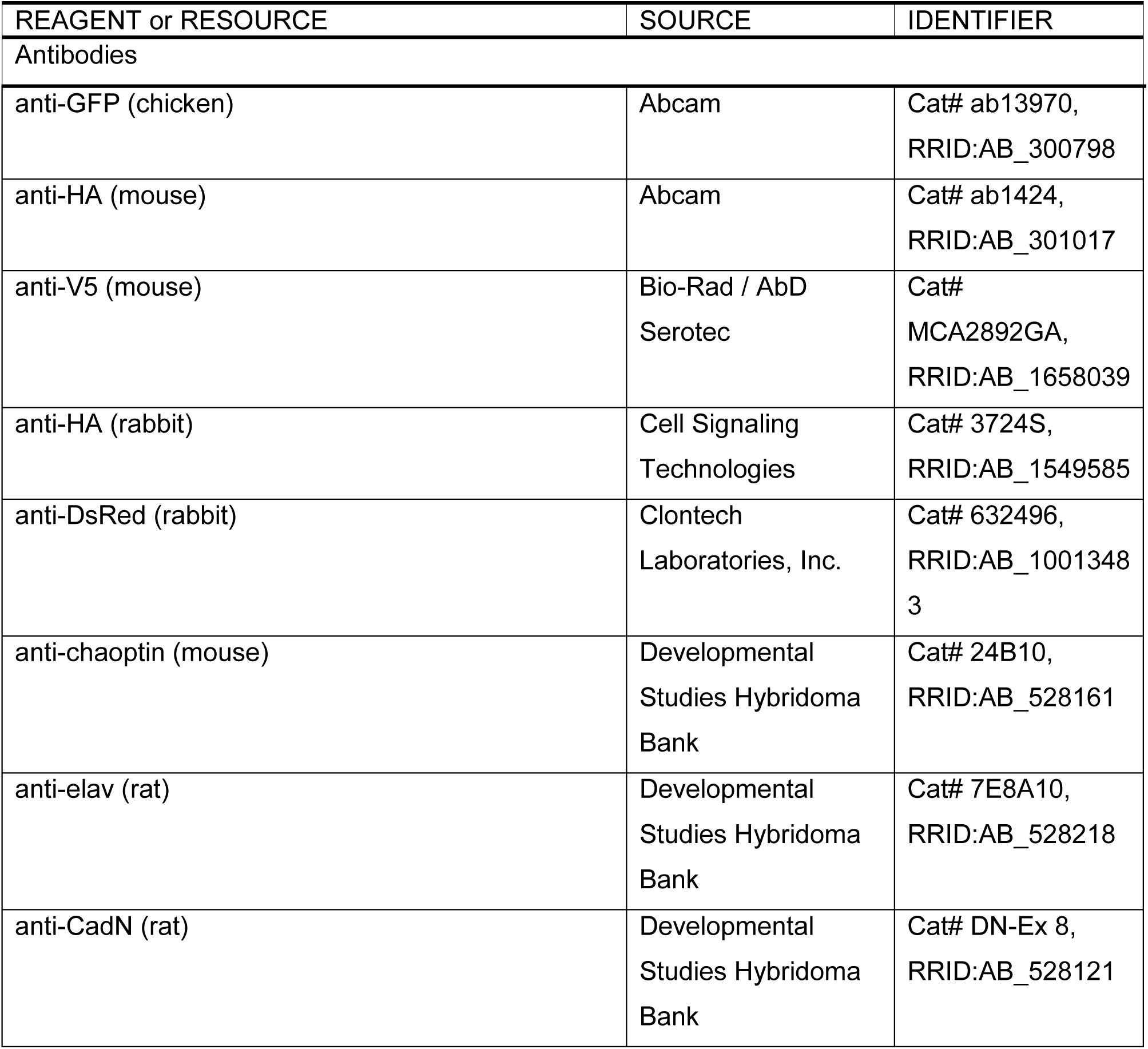
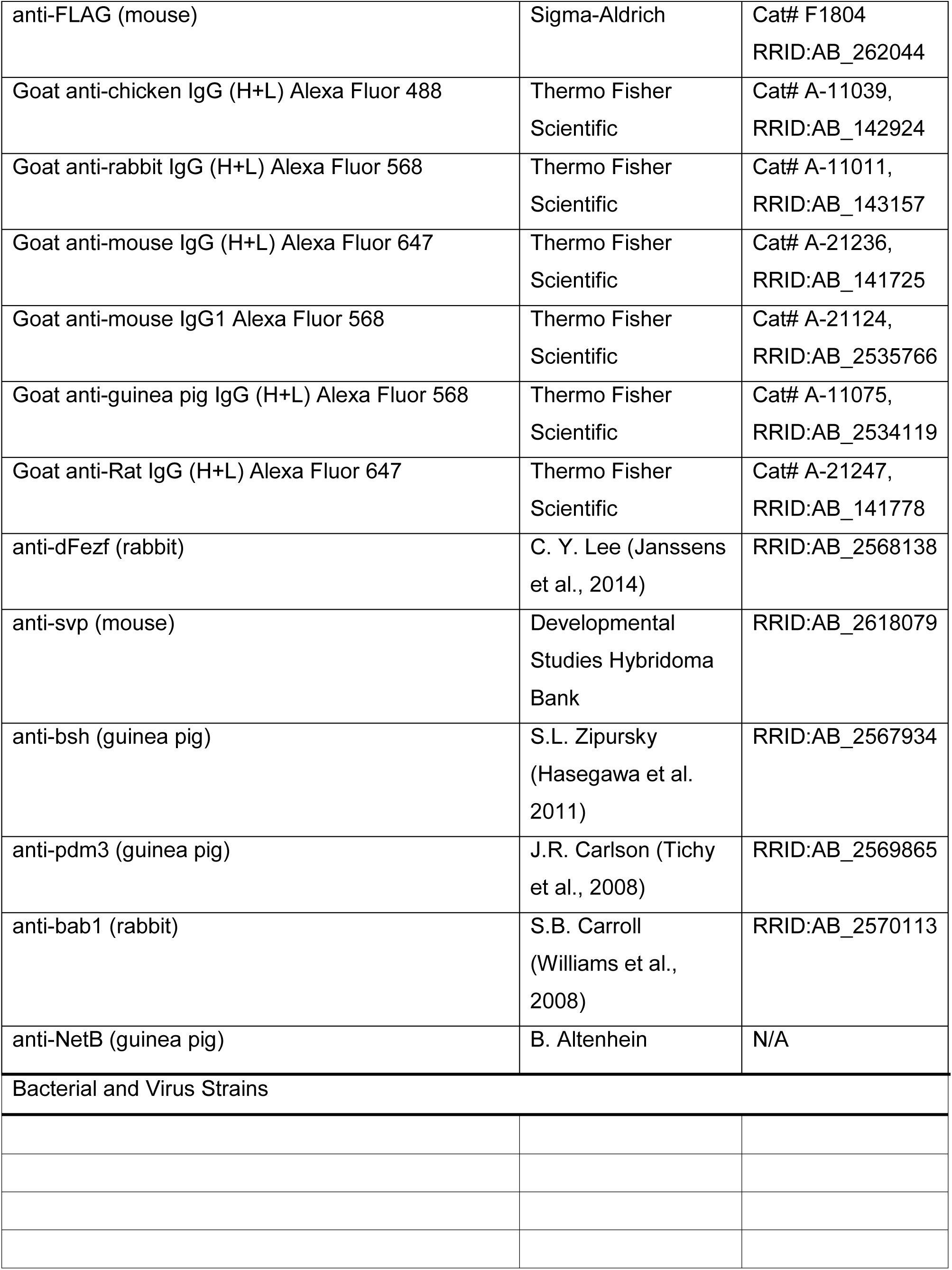
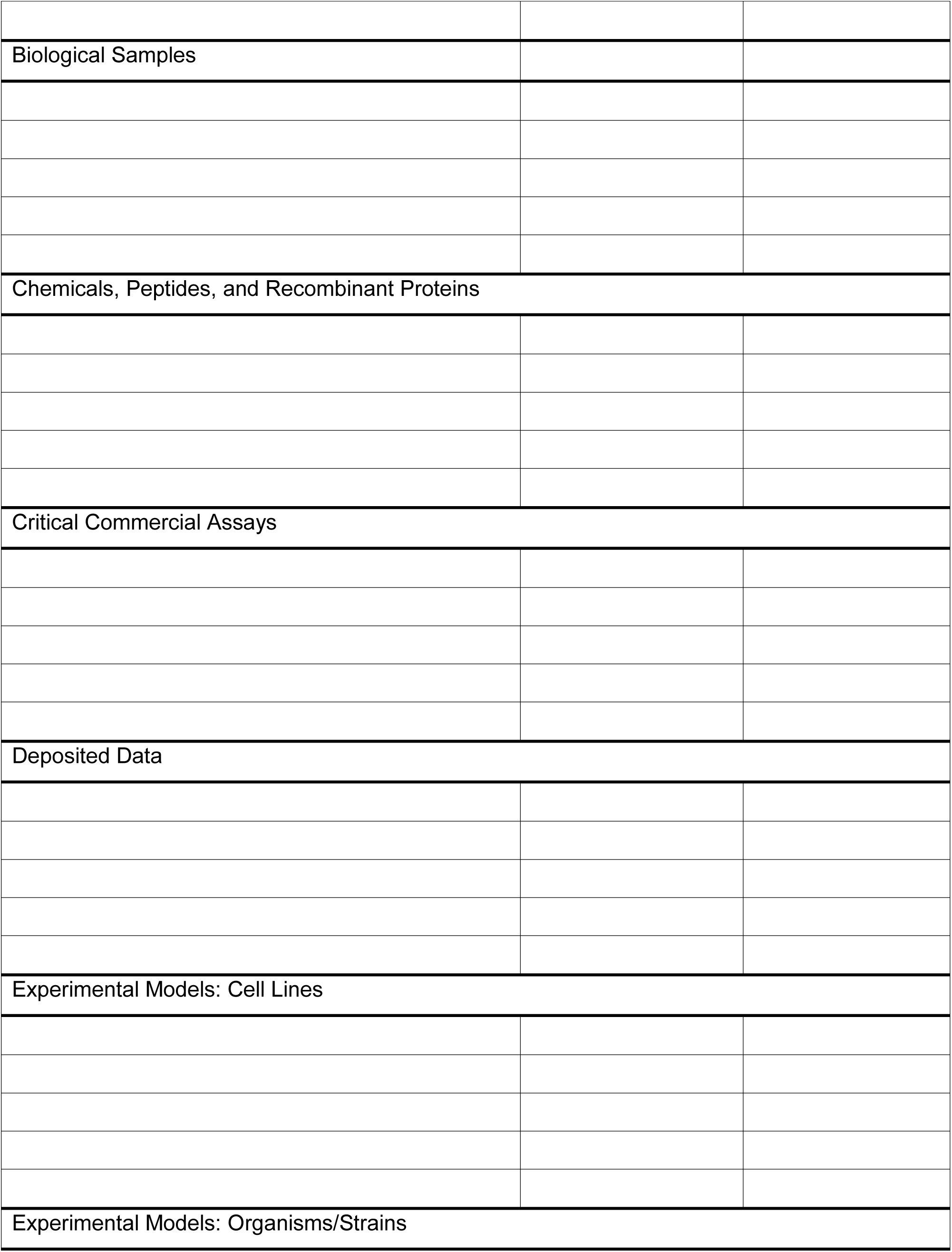
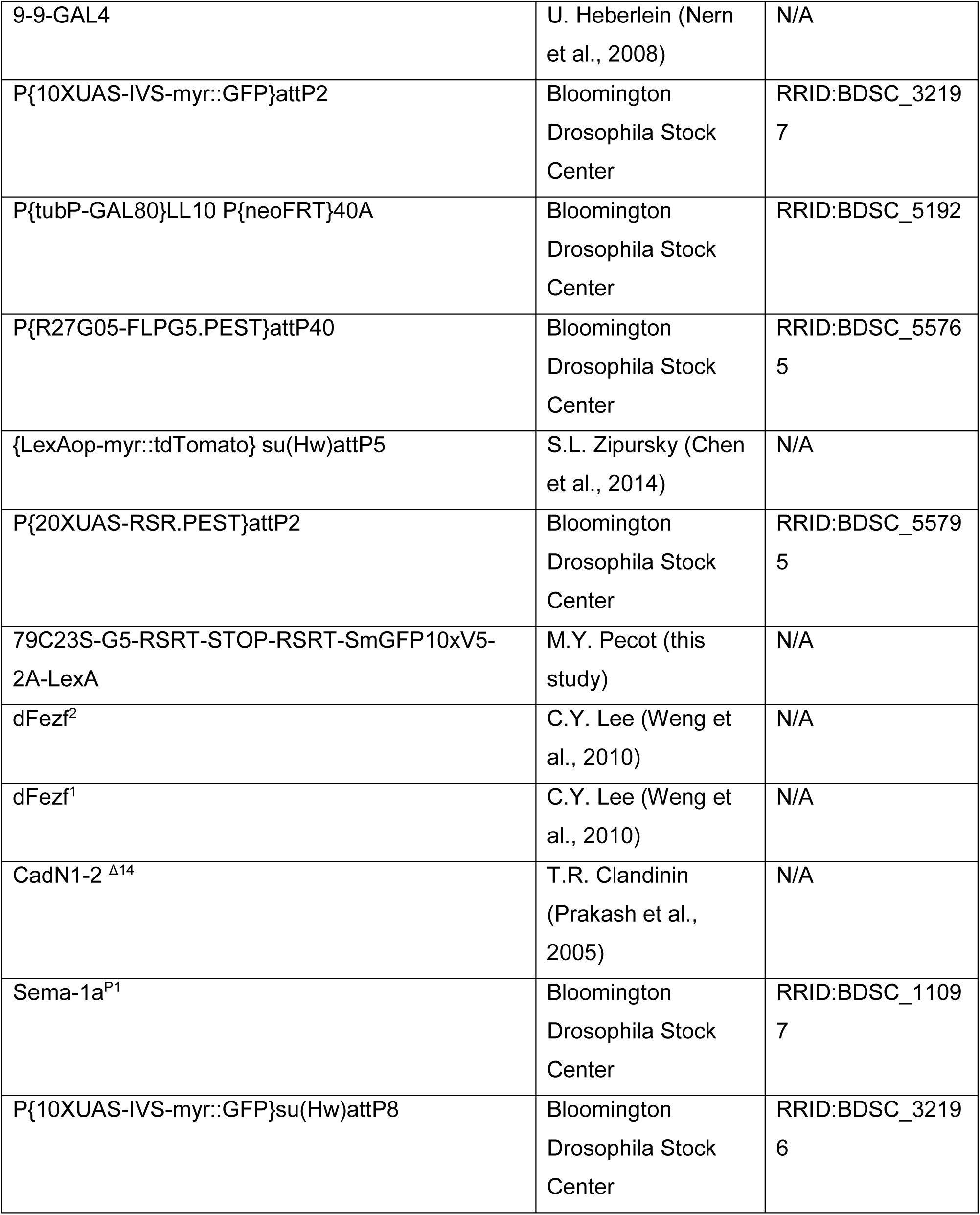
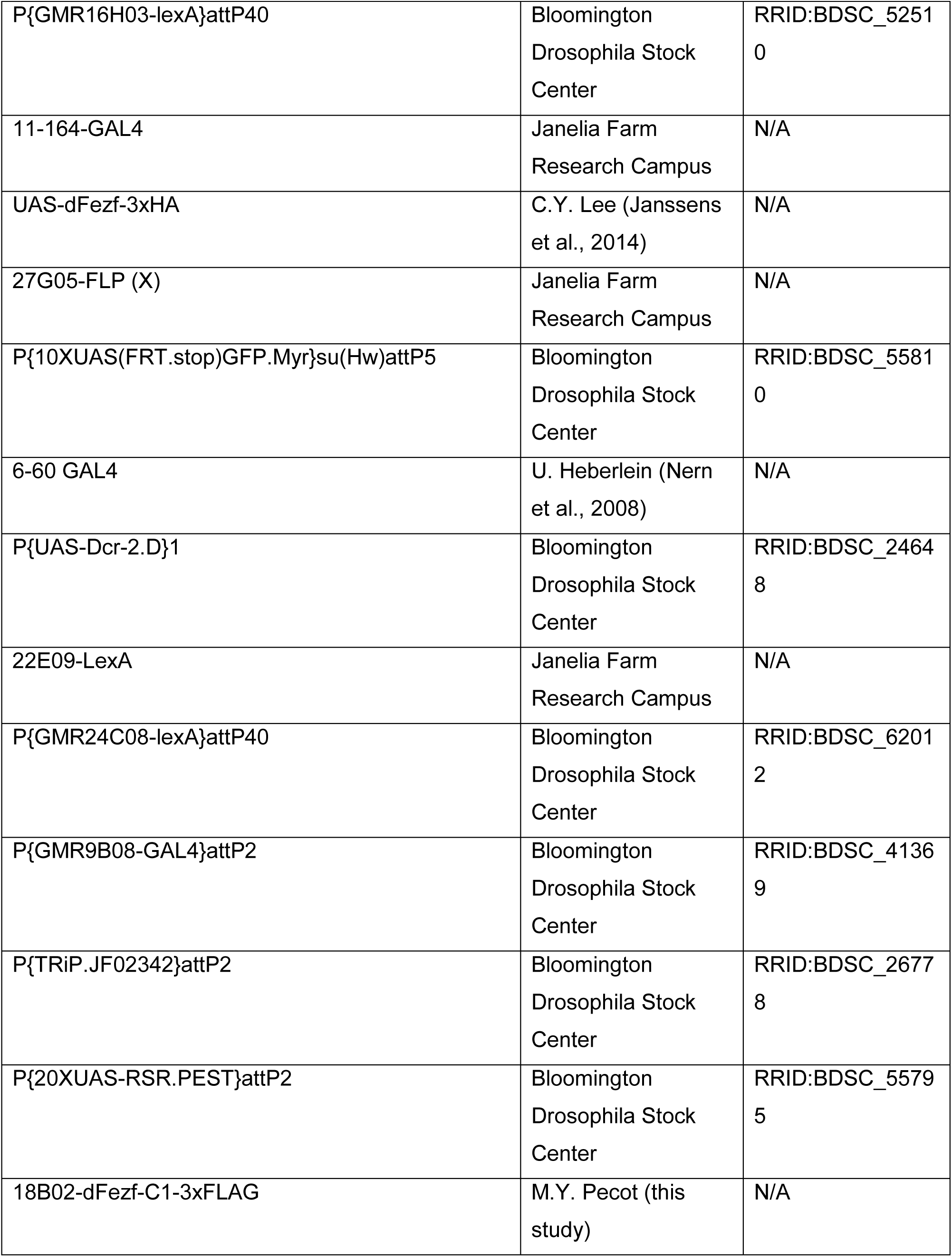
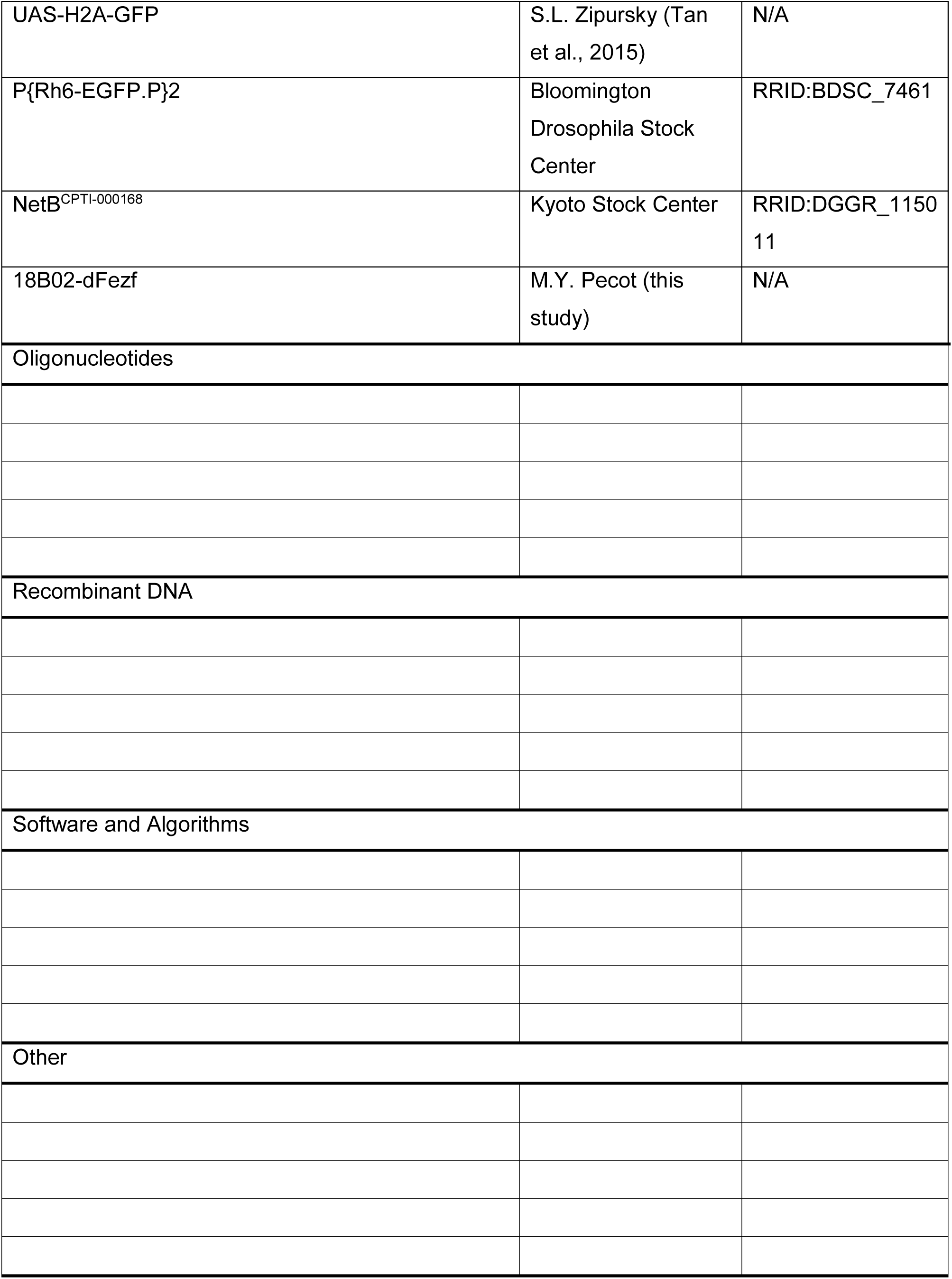

**Table S1.**
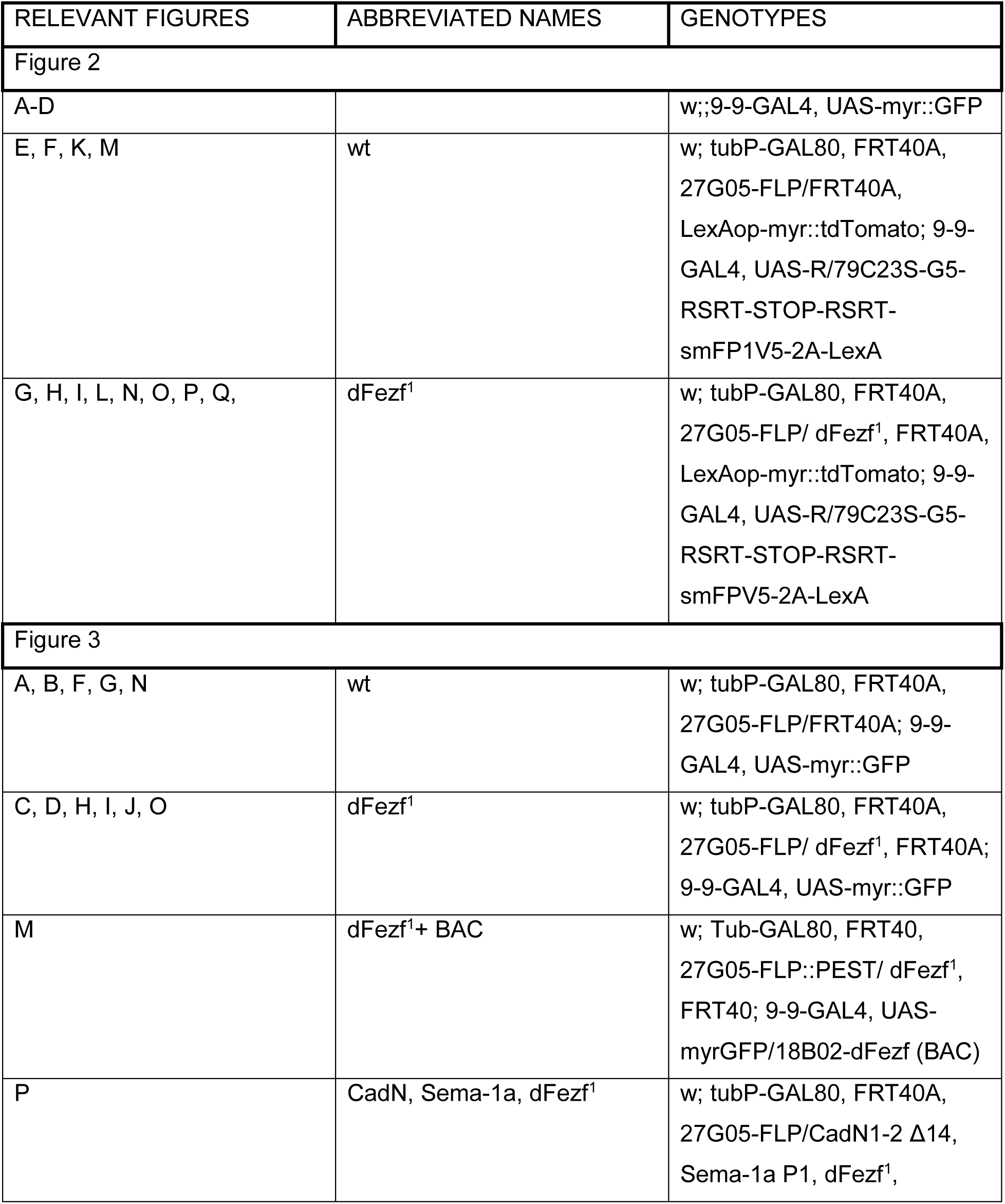
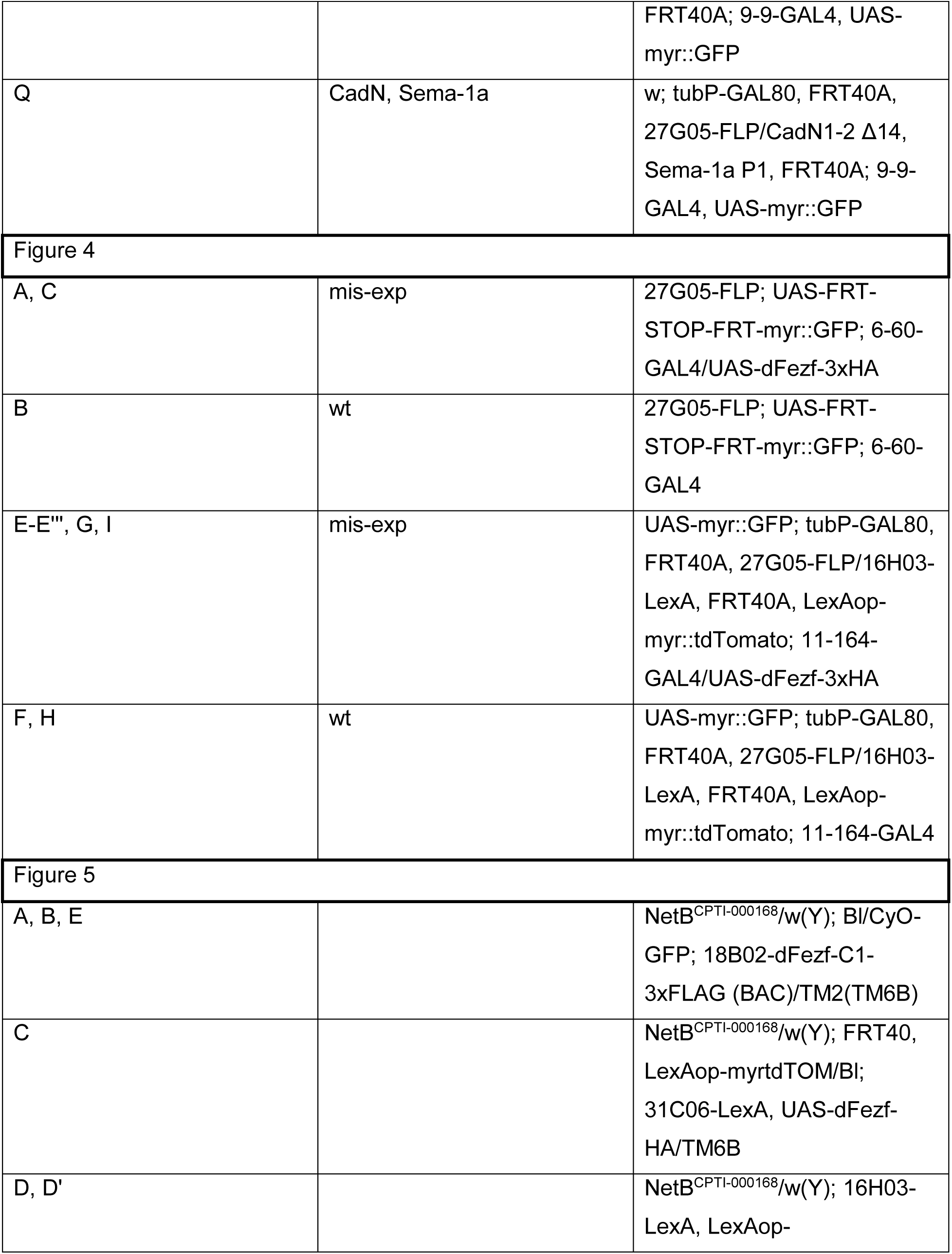
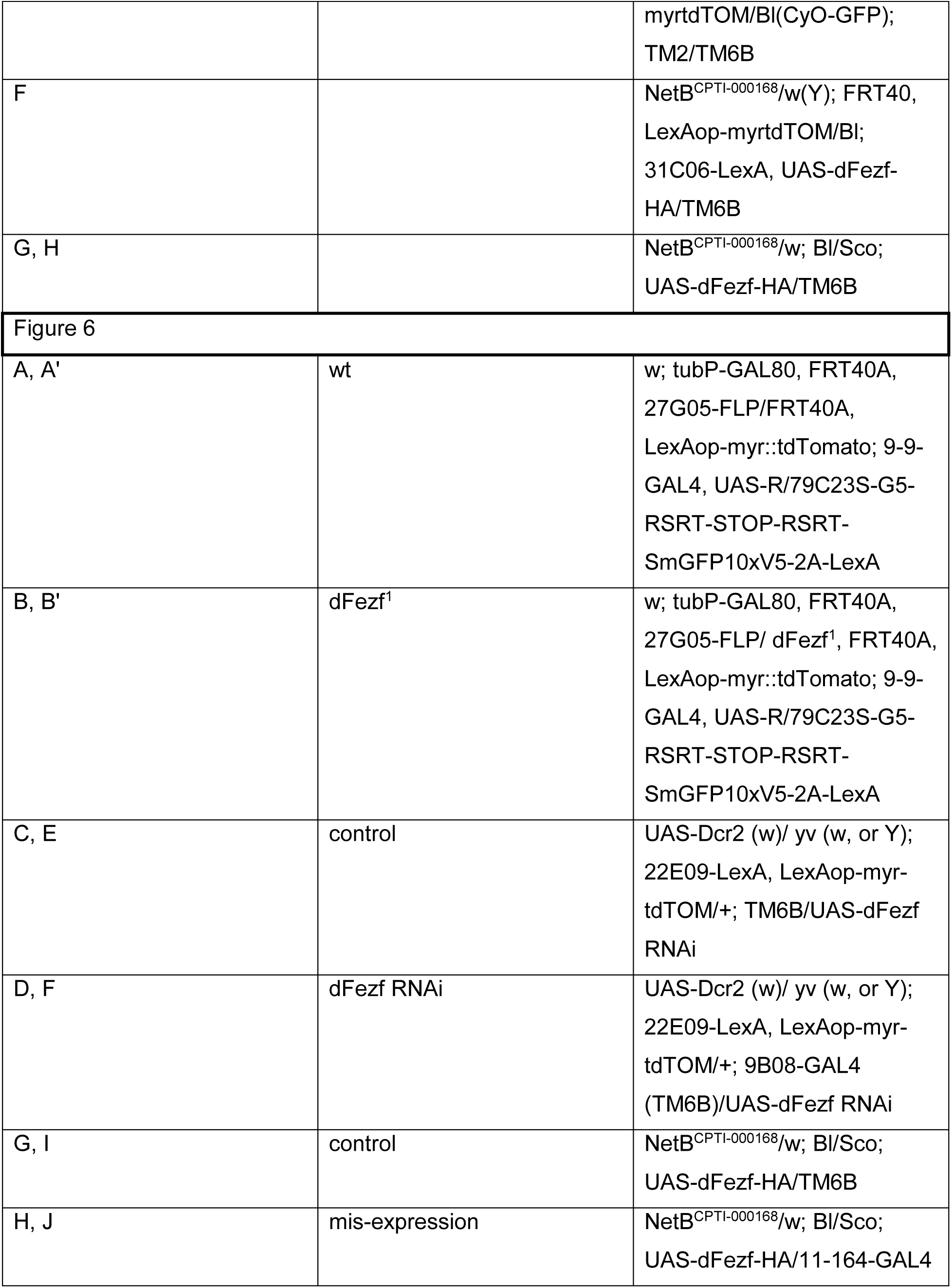
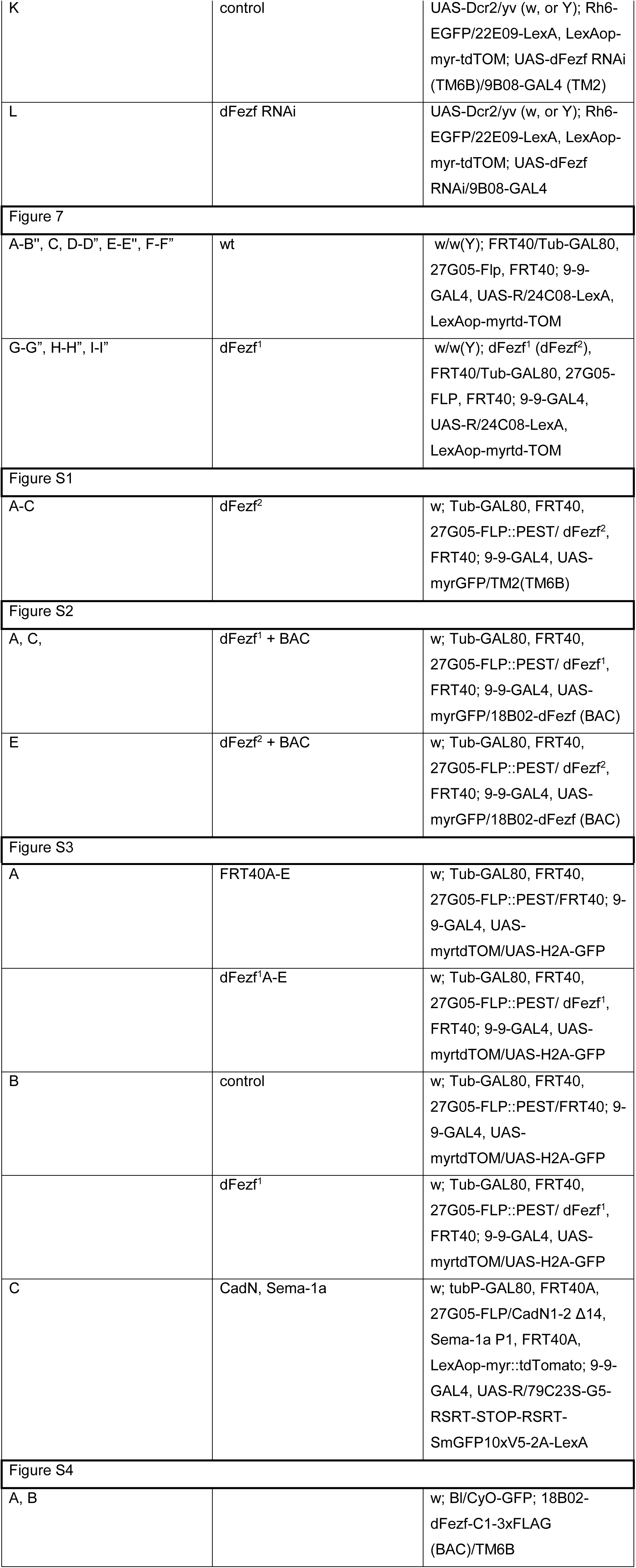
Full genotypes of flies in each experiment

| RELEVANT FIGURES | ABBREVIATED NAMES | GENOTYPES |
| --- | --- | --- |
| Figure 2 |  |  |
| A-D |  | w;;9-9-GAL4, UAS-myr::GFP |
| E, F, K, M | wt | w; tubP-GAL80, FRT40A, 27G05-FLP/FRT40A, LexAop-myr::tdTomato; 9-9-GAL4, UAS-R/79C23S-G5-RSRT-STOP-RSRT-smFP1V5-2A-LexA |
| G, H, I, L, N, O, P, Q, | dFezf<sup>1</sup> | w; tubP-GAL80, FRT40A, 27G05-FLP/ dFezf<sup>1</sup>, FRT40A, LexAop-myr::tdTomato; 9-9-GAL4, UAS-R/79C23S-G5-RSRT-STOP-RSRT-smFPV5-2A-LexA |
| Figure 3 |  |  |
| A, B, F, G, N | wt | w; tubP-GAL80, FRT40A, 27G05-FLP/FRT40A; 9-9-GAL4, UAS-myr::GFP |
| C, D, H, I, J, O | dFezf<sup>1</sup> | w; tubP-GAL80, FRT40A, 27G05-FLP/ dFezf<sup>1</sup>, FRT40A; 9-9-GAL4, UAS-myr::GFP |
| M | dFezf<sup>1</sup>+ BAC | w; Tub-GAL80, FRT40, 27G05-FLP::PEST/ dFezf<sup>1</sup>, FRT40; 9-9-GAL4, UAS-myrGFP/18B02-dFezf (BAC) |
| P | CadN, Sema-1a, dFezf<sup>1</sup> | w; tubP-GAL80, FRT40A, 27G05-FLP/CadN1-2 Δ14, Sema-1a P1, dFezf<sup>1</sup>, |
|  |  | FRT40A; 9-9-GAL4, UAS-myr::GFP |
| Q | CadN, Sema-1a | w; tubP-GAL80, FRT40A, 27G05-FLP/CadN1-2 Δ14, Sema-1a P1, FRT40A; 9-9-GAL4, UAS-myr::GFP |
| Figure 4 |  |  |
| A, C | mis-exp | 27G05-FLP; UAS-FRT-STOP-FRT-myr::GFP; 6-60-GAL4/UAS-dFefz-3xHA |
| B | wt | 27G05-FLP; UAS-FRT-STOP-FRT-myr::GFP; 6-60-GAL4 |
| E-E''', G, I | mis-exp | UAS-myr::GFP; tubP-GAL80, FRT40A, 27G05-FLP/16H03-LexA, FRT40A, LexAop-myr::tdTomato; 11-164-GAL4/UAS-dFefz-3xHA |
| F, H | wt | UAS-myr::GFP; tubP-GAL80, FRT40A, 27G05-FLP/16H03-LexA, FRT40A, LexAop-myr::tdTomato; 11-164-GAL4 |
| Figure 5 |  |  |
| A, B, E |  | NetB<sup>CPTI-000168</sup>/w(Y); BI/CyO-GFP; 18B02-dFefz-C1-3xFLAG (BAC)/TM2(TM6B) |
| C |  | NetB<sup>CPTI-000168</sup>/w(Y); FRT40, LexAop-myrtdTOM/BI; 31C06-LexA, UAS-dFefz-HA/TM6B |
| D, D' |  | NetB<sup>CPTI-000168</sup>/w(Y); 16H03-LexA, LexAop- |
|  |  | myrtdTOM/BI(CyO-GFP);<br>TM2/TM6B |
| F |  | NetB<sup>CPTI-000168</sup>/w(Y); FRT40,<br>LexAop-myrtdTOM/BI;<br>31C06-LexA, UAS-dFezf-HA/TM6B |
| G, H |  | NetB<sup>CPTI-000168</sup>/w; BI/Sco;<br>UAS-dFezf-HA/TM6B |
| Figure 6 |  |  |
| A, A' | wt | w; tubP-GAL80, FRT40A,<br>27G05-FLP/FRT40A,<br>LexAop-myr::tdTomato; 9-9-GAL4, UAS-R/79C23S-G5-RSRT-STOP-RSRT-SmGFP10xV5-2A-LexA |
| B, B' | dFezf<sup>1</sup> | w; tubP-GAL80, FRT40A,<br>27G05-FLP/ dFezf<sup>1</sup>, FRT40A,<br>LexAop-myr::tdTomato; 9-9-GAL4, UAS-R/79C23S-G5-RSRT-STOP-RSRT-SmGFP10xV5-2A-LexA |
| C, E | control | UAS-Dcr2 (w)/ yv (w, or Y);<br>22E09-LexA, LexAop-myr-tdTOM/+; TM6B/UAS-dFezf<br>RNAi |
| D, F | dFezf RNAi | UAS-Dcr2 (w)/ yv (w, or Y);<br>22E09-LexA, LexAop-myr-tdTOM/+; 9B08-GAL4<br>(TM6B)/UAS-dFezf RNAi |
| G, I | control | NetB<sup>CPTI-000168</sup>/w; BI/Sco;<br>UAS-dFezf-HA/TM6B |
| H, J | mis-expression | NetB<sup>CPTI-000168</sup>/w; BI/Sco;<br>UAS-dFezf-HA/11-164-GAL4 |
| K | control | UAS-Dcr2/yv (w, or Y); Rh6-EGFP/22E09-LexA, LexAop-myr-tdTOM; UAS-dFezf RNAi (TM6B)/9B08-GAL4 (TM2) |
| L | dFezf RNAi | UAS-Dcr2/yv (w, or Y); Rh6-EGFP/22E09-LexA, LexAop-myr-tdTOM; UAS-dFezf RNAi/9B08-GAL4 |
| Figure 7 |  |  |
| A-B", C, D-D", E-E", F-F" | wt | w/w(Y); FRT40/Tub-GAL80, 27G05-Flp, FRT40; 9-9-GAL4, UAS-R/24C08-LexA, LexAop-myrtd-TOM |
| G-G", H-H", I-I" | dFezf<sup>1</sup> | w/w(Y); dFezf<sup>1</sup> (dFezf<sup>2</sup>), FRT40/Tub-GAL80, 27G05-FLP, FRT40; 9-9-GAL4, UAS-R/24C08-LexA, LexAop-myrtd-TOM |
| Figure S1 |  |  |
| A-C | dFezf<sup>2</sup> | w; Tub-GAL80, FRT40, 27G05-FLP::PEST/ dFezf<sup>2</sup>, FRT40; 9-9-GAL4, UAS-myrGFP/TM2(TM6B) |
| Figure S2 |  |  |
| A, C, | dFezf<sup>1</sup> + BAC | w; Tub-GAL80, FRT40, 27G05-FLP::PEST/ dFezf<sup>1</sup>, FRT40; 9-9-GAL4, UAS-myrGFP/18B02-dFezf (BAC) |
| E | dFezf<sup>2</sup> + BAC | w; Tub-GAL80, FRT40, 27G05-FLP::PEST/ dFezf<sup>2</sup>, FRT40; 9-9-GAL4, UAS-myrGFP/18B02-dFezf (BAC) |
| Figure S3 |  |  |
| A | FRT40A-E | w; Tub-GAL80, FRT40, 27G05-FLP::PEST/FRT40; 9-9-GAL4, UAS-myrtdTOM/UAS-H2A-GFP |
|  | dFezf<sup>1</sup>A-E | w; Tub-GAL80, FRT40, 27G05-FLP::PEST/ dFezf<sup>1</sup>, FRT40; 9-9-GAL4, UAS-myrtdTOM/UAS-H2A-GFP |
| B | control | w; Tub-GAL80, FRT40, 27G05-FLP::PEST/FRT40; 9-9-GAL4, UAS-myrtdTOM/UAS-H2A-GFP |
|  | dFezf<sup>1</sup> | w; Tub-GAL80, FRT40, 27G05-FLP::PEST/ dFezf<sup>1</sup>, FRT40; 9-9-GAL4, UAS-myrtdTOM/UAS-H2A-GFP |
| C | CadN, Sema-1a | w; tubP-GAL80, FRT40A, 27G05-FLP/CadN1-2 Δ14, Sema-1a P1, FRT40A, LexAop-myr::tdTomato; 9-9-GAL4, UAS-R/79C23S-G5-RSRT-STOP-RSRT-SmGFP10xV5-2A-LexA |
| Figure S4 |  |  |
| A, B |  | w; Bl/CyO-GFP; 18B02-dFezf-C1-3xFLAG (BAC)/TM6B |

